# Tracing India’s Canine Heritage through SNP-Based Haplotype Identification

**DOI:** 10.1101/2024.01.02.573915

**Authors:** Dapinder Singh, Shashi Kant Mahajan, Neeraj Kashyap, Chandra Sekhar Mukhopadhyay

**Affiliations:** Department of Bioinformatics, College of Animal Biotechnology, Guru Angad Dev Veterinary and Animal Sciences University, Ludhiana, Punjab, India; Department of Veterinary Surgery & Radiology, College of Veterinary Sciences, Guru Angad Dev Veterinary and Animal Sciences University, Ludhiana, Punjab, India; Department of Animal Genetics & Breeding, College of Veterinary Science, Guru Angad Dev Veterinary and Animal Sciences University, Ludhiana, Punjab, India

**Keywords:** Population structure, Haplotype identification, R-coding, Dogs, SNPs

## Abstract

Dog breeds/germplasm in India is mostly unexplored and the population structure of owned dogs has not been studied at all. The current study was designed to determine the haplotypes and explore the population structure among divergent breeds of dogs using genome-wide distributed SNPs, followed by validation of selected haplotypes through PCR-sequencing. The research employed custom ddRAD-GBS sequencing carried out using Illumina 150bp paired-end sequencing of 50 dog samples generated 2,18,433 high-quality SNPs meeting the screening criteria. The data was further analyzed for population structure assessment and haplotype identification using bash, and R-environment. Subsequently, three haplotypes (on AFAP1, CELSR1, and GBGT1 genes) were selected (based on SNP density and haplotype length) for validation via PCR followed by paired-end Sanger sequencing in seven different dog breeds (n=21). The results revealed notable connections between dog breeds from Punjab and Haryana, while the affiliation with Karnataka was found to be less pronounced. The sequencing results indicated that CELSR1 and GBGT1 genes contained SNPs, with the AFAP1 gene lacking SNPs. The results also provided insights into the molecular-level population structure, SNPs, and haplotypes of diverse dog breeds reared in India. This SNP variation could be used for molecular characterization of indigenous dogs.

## Introduction

Dog is one of the first animals to be domesticated and is the most well-known pet in human history. Modern domesticated dogs have been selected artificially for over 14,000 years as pets or guarding animals or hunting companions. According to the American Kennel Club (AKC), 199 dog breeds are recognized worldwide out of a total of 340 dog breeds known to exist worldwide that exhibit significant variations in morpho-physiological and behavioural characteristics (https://www.akc.org/dog-breeds/). Dog populations have developed into hundreds of breeds over thousands of years, each of which is distinguished by traits that give a distinctive range of polymorphism. SNPs can be one such cause that has produced an incredible diversity of dog breeds that have developed much more recently (Ali *et al*. 2020). A single nucleotide polymorphism (SNP) is a DNA locus or site where individuals within a species differ from one another. They are the most abundant type of polymorphism in animal genomes. The SNP data is widely used for determining the population structure of various plant and animal species (Liu *et al*. 2022).

Haplotypes are the set of polymorphic alleles or SNPs that co-occur on a chromosome and tend to be inherited together. The patterns of haplotype diversity and sharing in and among populations closely parallel those described for SNPs. Haplotype identification is important to study the genetic diversity and population structure of different dog breeds. Furthermore, the number of haplotypes for a gene is strongly correlated with the number of SNPs (Vychodilova *et al*. 2018).

Population structure which is defined as the occurrence of a systematic differentiation in allele frequencies between sub-populations as a result of non-random mating between individuals (Ubbens *et al*. 2022), is useful in determining the evolutionary history of the breeds.

The primary method commonly employed to explore the genetic makeup of populations is illustrated through principal components analysis (PCA), as described by van Waaij *et al*. (2023). In this approach, the matrix under scrutiny contains information about how genetically similar pairs of individuals are. The principal components (PCs) of this matrix signify the most significant directions within the sample space, effectively capturing the main patterns of genetic resemblance that are observed.

Use of high-throughput sequencing technology to discover a large number of SNPs has proved to be not time-consuming and cost-effective (Eriksson *et al*. 2020). The usage of the genotyping-by-sequencing (GBS) technique has increased dramatically with the rapid development of next-generation (NGS) technologies used in plant breeding programs (Elshire *et al*. 2011). GBS enormously reduces the complexity of large genomes of species by choosing appropriate restriction enzymes (REs). Poland and colleagues have developed a GBS protocol using two REs (*PstI*/*MspI*), which can reduce complexity to a greater extent and achieve a more unified sequencing library than a one-enzyme protocol (Poland *et al*. 2012).

The major goal of this study is to describe the population structure in the native dog breeds by utilizing genome-wide SNPs and to identify haplotypes from them. Research reports on the molecular characterization of dogs are very limited. Here we use a genomic approach including 2,18,433 genome-wide SNPs obtained from ddRAD-GBS. The number of canine SNPs is still unknown, and nearly all of them are still unidentified. However, there is no information is available about the molecular characterization of divergent breeds of owned dogs. Identification of haplotypes along with analysis of population structure remains unknown in the owned dog breeds reared in India. It has been hypothesized that the population structure and haplotype can be assessed from the genome-wide SNPs of the commonly available dog breeds in India. Thus, the present study aimed to elucidate the haplotypes and explore the population structure among divergent breeds of dogs in India and validate selected haplotypes in Labrador retriever and German shepherd breeds through PCR-sequencing and analysis of SNPs.

## Materials and methods

### Sample collection and DNA extraction for GBS data

The Genotyping-by-sequencing (GBS) data were already available in the lab. As sample collection, DNA extraction and customized GBS subjected to Illumina sequencing have already been completed; the brief overview of the samples that were gathered is given in Table 1.

**Table 1.**
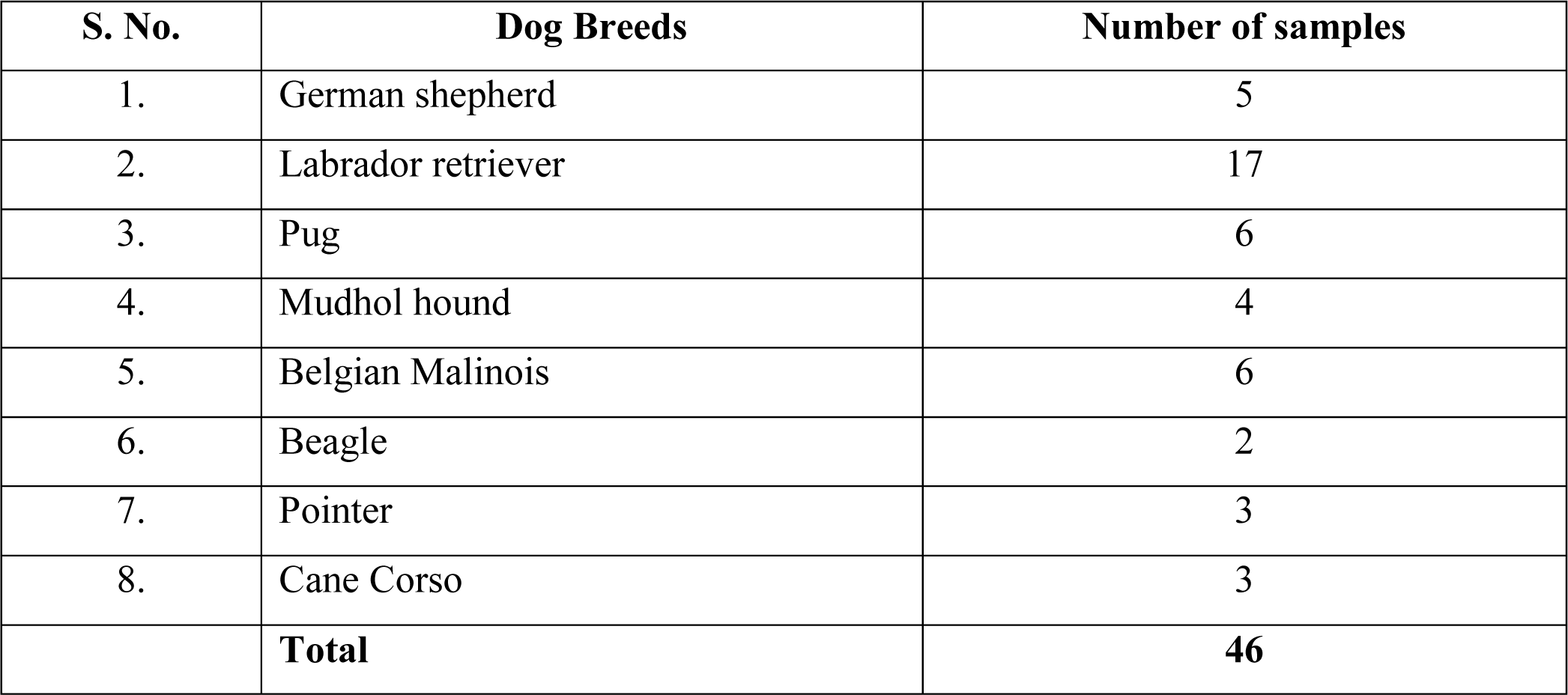
Number of samples by breed used in current study.

| S. No. | Dog Breeds | Number of samples |
| --- | --- | --- |
| 1. | German shepherd | 5 |
| 2. | Labrador retriever | 17 |
| 3. | Pug | 6 |
| 4. | Mudhol hound | 4 |
| 5. | Belgian Malinois | 6 |
| 6. | Beagle | 2 |
| 7. | Pointer | 3 |
| 8. | Cane Corso | 3 |
|  | **Total** | **46** |

Whole-blood samples (2 ml) were collected from fifty canines belonging to eight different breeds from different states of India viz. Punjab (n = 38), Haryana (n = 6) and Karnataka (n = 6). The samples were drawn from cephalic vein using blood collection tubes containing EDTA as an anticoagulant. Genomic DNA extractions were performed following phenol: chloroform: isoamyl alcohol method (Green & Sambrook, 2018). The DNA samples were quantified using the Thermo Scientific™ NanoDrop 2000 spectrophotometer to assess the purity of the extracted DNA. Furthermore, the DNA samples were checked for molecular quality by running them through a 0.8% agarose gel against a 1 kilobase (kb) DNA ladder marker. All the samples were sequenced using the GBS platform, for which an initial quality control (QC) report of 50 samples was received but after genotyping-by-sequencing (GBS), quality control (QC) report for only 46 samples were received as 4 samples (viz. German Shepherd = 1, Labrador = 1 and Mudhol Hound = 2) were found insufficient for sequencing.

### ddRAD-Genotyping by Sequencing

The PCR products were then sent to Novogene Lifesciences Pvt. Ltd., Mumbai (https://en.novogene.com/), where the GBS approach was carried out following the GBS protocol (Peterson *et al*. 2012) along with mapping the SNPs to the reference dog genome (https://www.ncbi.nlm.nih.gov/assembly/GCF_014441545.1/) BioProject - PRJNA843534; Biosamples - SAMN35074275 and SRA-Accessions - SRR24547516. The GBS DNA library was prepared using the ddRAD approach for each sample by digesting them with two restriction enzymes (*MseI* & *HaeIII_MspI*) and the resulting fragments were joined by two barcoded adapters that either had a compatible sticky end with universal Illumina P5 or P7 sequences or with the primary digestion enzyme. After the sequencing evaluation summary, the raw data for 46 samples were sequenced as the four samples failed the initial quality control. Using BWA (Li & Durbin, 2009) software, the effective sequencing data was aligned with the reference sequence (parameters: mem -t 4 -k 32 -M), and the mapping rate and coverage were calculated following the alignment results. SAMTOOLS (Li *et al*. 2009) was used to handle the BAM files.

### SNP calling

SAMtools (Li *et al*. 2009) was used to detect the individual SNP variation by using (mpileup -m 2 -F 0.002 -d 1000) parameter. To lower the error rate in SNP detection the following parameters were utilized: (a) there should be more than 4 support reads for each SNP, and (b) each SNP should have a mapping quality (MQ) of greater than 20.

### Raw data cleaning

The raw data is refined by verifying accession numbers with chromosome details and removing entries without this information. The dataset is then split into 13 trios using R-coding for Mendelian error assessment. Errors like tri-allelic genotypes separated by a tab are removed. Trios’ parental genotypes are merged and evaluated for Mendelian errors. Non-Mendelian SNPs are filtered based on segregation distortion, deviating from Mendelian inheritance. SNPs file is then converted to genotypes files, maintaining data but in a more manageable format for analysis. Genotype files organize data in a tabular structure for individuals and SNPs, aiding analysis. This conversion facilitates Minor Allele Frequency (MAF) calculation.

### Investigating canine population structure using R-programming

Filtered VCF files were analyzed in R v4.3.1 using vcfR, poppr, ape, and RColorBrewer packages. The “dist()” function generated a pairwise genetic distance matrix, indicating genetic distances between all individuals. Genetic relationships were assessed with a UPGMA tree based on samples from Punjab, Haryana, and Karnataka (46 samples, cutoff 50). Genetic distance was calculated using bitwise.dist() function from the genlight object. Tree tips were color-coded by the population from which the samples originated. For population-wide relationships and multilocus genotypes (MLG), a minimum spanning network (MSN) was constructed using a genlight object and distance matrix. Allelic differences were calculated by bitwise.dist() from poppr.

Principal Component Analysis (PCA) converts SNP data into uncorrelated principal components, summarizing sample variation. Using the glPCA() function on the genlight object the principal components were generated. These are incorporated into a new object with added population values for coloration according to their state-wise location. Visualization is achieved via ggplot2. Discriminant Analysis of Principal Components (DAPC) determines an optimal number of genetically differentiated clusters. DAPC uses a genlight object with preset populations, utilizing PCA results for sample assignment. In this case, the objective is to assign each sample to a population based on the results of the PCA. Therefore, the same parameters used in the PCA (n.pca=3 and n.da=2) were used to reconstruct the DAPC. ggplot2 and tidyr transform data for scatter plot visualization.

For clearer interpretation, we reconstructed the plots using ggplot2 and grouped samples by their locations. We extracted DAPC-calculated population assignments and formed a new data frame with original population labels and sample names. Using pivot_longer from tidyr, we reshaped the data frame to meet ggplot2’s structure requirements. Columns were renamed for familiarity. The restructured dapc.results data frame was plotted using ggplot2. Samples are on the X-axis, membership probabilities are on the Y-axis, and fill colour shows original populations. Each facet in the bar plot depicts the initial population assignment, giving an organized view contrasting membership probabilities with original populations.

### Haplotype identification

Haplotype-based analysis, when compared to the utilization of individual SNPs, can effectively lower the occurrence of false discoveries due to the reduced number of association tests conducted (Hamblin & Jannink, 2011). For any specific segment of chromosomal DNA, an individual can possess two haplotypes, while at the population level, there can be multiple haplotypes present for that same segment of chromosomal DNA. The haplotype identification from the SNP genotype data was performed using an R-programming environment in the Ubuntu Linux terminal. The following criteria were followed for the identification of the haplotypes: (a) distance between SNP – less than 80kb, (b) haplotype length – up to 150kb, and (c) at least 4 SNPs should be present in one haplotype.

### Validation of haplotypes

Three haplotypes were chosen for the validation studies considering SNP density and MAF values of 0.05 or 5%. To identify associated genes with the selected haplotypes, NCBI (https://www.ncbi.nlm.nih.gov/) was queried for the respective RefSeq IDs. Three genes namely *AFAP1*, *GBGT1* and *CELSR1* were selected and the primers for these genes were self-designed using Primer-BLAST (https://www.ncbi.nlm.nih.gov/tools/primer-blast/). The details of the primers are given in Table 2

**Table 2.**
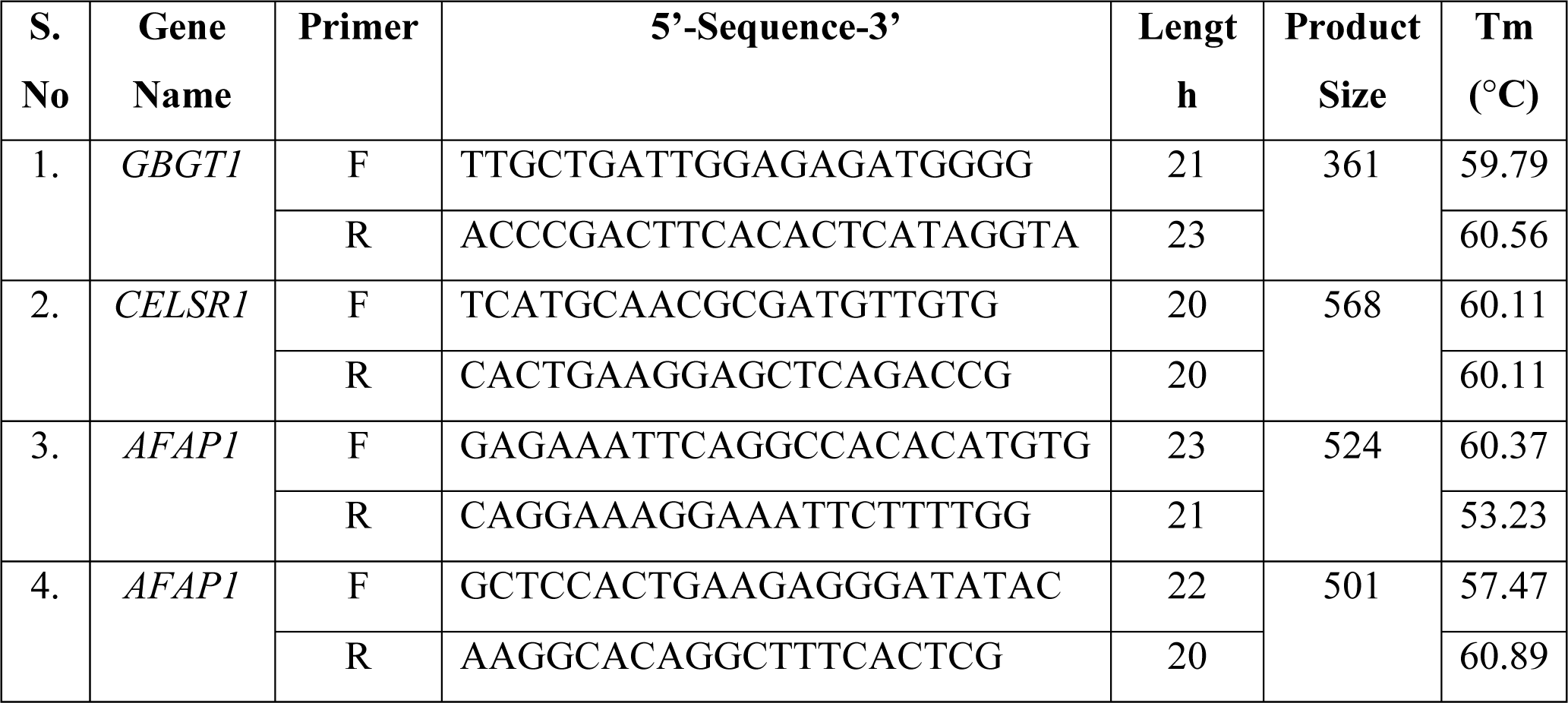
Primer details of target gene amplification.

### Sample collection for haplotype and SNP validation

Blood sample collection from dogs was authorized by the Institutional Academic Ethical Committee (IAEC), as per Memo No.: IAEC/2023/134-55 Dated 22/05/2023 and Memo No.: IAEC/2023/176-186 Dated 14/09/2023. Samples were aseptically collected from Guru Angad Dev Veterinary and Animal Sciences University (GADVASU) Multispecialty clinics. Seven different breeds were sampled using a 21G needle (DISPO VAN^TM^) from the cephalic vein (Table 3). Healthy dog samples were collected regardless of age and sex into 15ml sterile tubes with anticoagulant (EDTA). Samples were stored at −20°C until analysis.

**Table 3.**
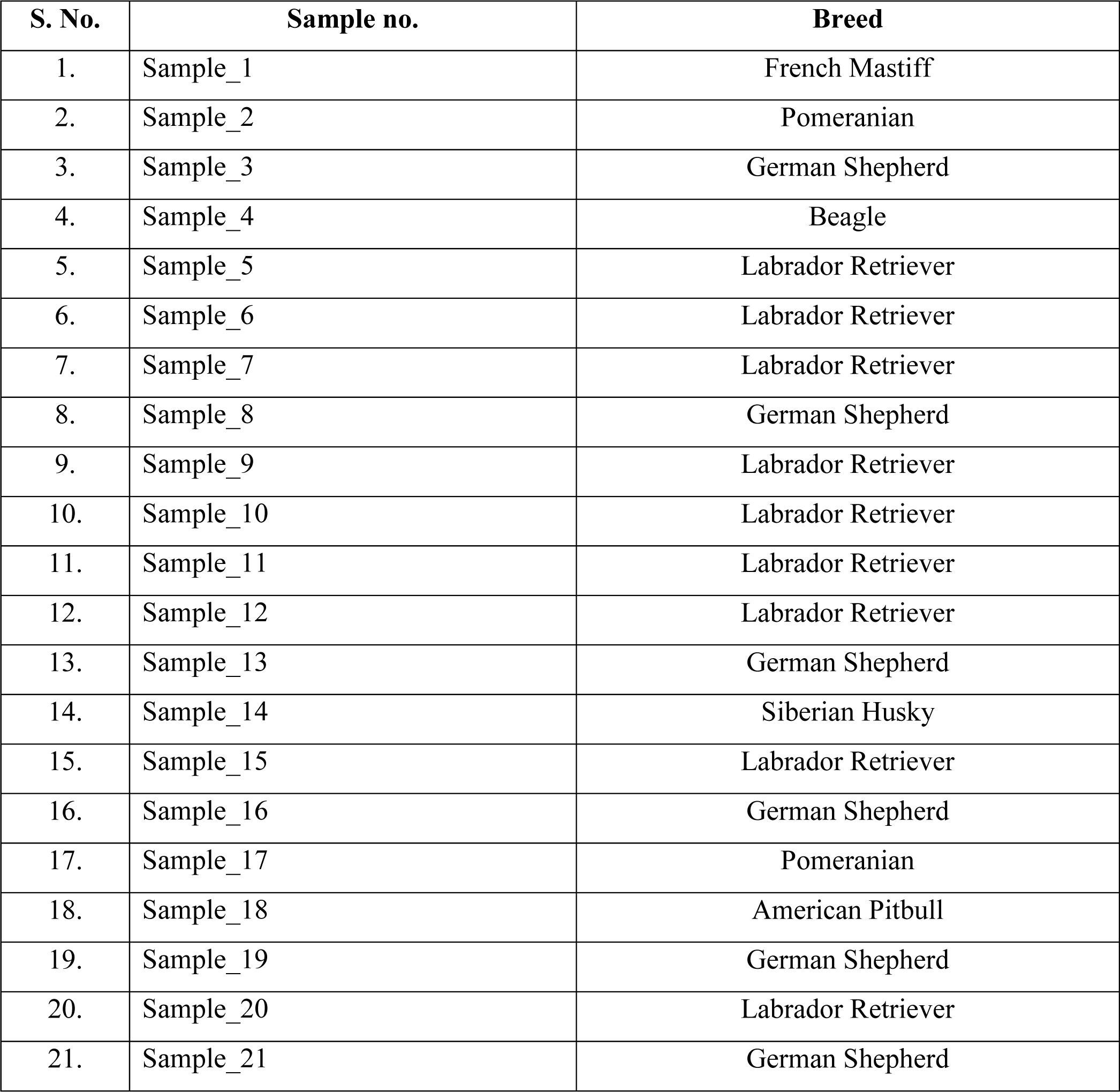
Blood samples collected from seven different dog breeds for validation studies.

### Sequencing of PCR products for identification of SNPs

PCR products of the *GBGT1*, *CELSR1* and *AFAP1* genes were subjected to DNA sequencing. The purified PCR products underwent bidirectional sequencing (forward and reverse) at GeneSpec Pvt. Ltd., Kerala. Sequencing data were then visualized using FinchTV software (https://digitalworldbiology.com/FinchTV). The resulting FASTA sequences were aligned against reference chromosome sequences of respective genes using Clustal Omega (https://www.ebi.ac.uk/Tools/msa/clustalo/). For SNP genotyping, all chromatograms were manually screened at each detected SNP position across the twelve dogs.

### Associating SNPs with important traits in dogs through pathway and enrichment analysis

*In-silico* analysis of three genes involved pathway analysis through the DAVID database, a bioinformatics tool for gene functional classification. DAVID aids gene categorization using features like Annotation Tool, GoChart, KeggCharts, and Domain Charts (Sherman *et al*. 2022). For gene ontology, PANTHER 17.0 was employed, separately analyzing biological processes, cellular components, molecular functions, and protein classes. GeneCards database is used to study gene functions and pathways associated with three selected genes in humans.

### Evolutionary analysis for haplotype sequences using MEGA 11

MEGA (Molecular Evolutionary Genetics Analysis) is specialized software for molecular evolution analysis and phylogenetic tree construction. It’s designed for comparing homologous gene sequences within or across species, emphasizing evolutionary relationships and DNA/protein evolution patterns. MEGA provides tools for statistical analysis, enabling pairwise genetic distance calculation for insights into sequence divergence and similarity (Tamura *et al*. 2021). Using MEGA 11, we aligned the twelve FASTA sequences (*AFAP1*, *CELSR1*, and *GBGT1* genes) individually. The resulting MEGA-format files were used for constructing phylogenetic trees and computing pairwise distance matrices to determine evolutionary distances.

## Results

### Population structure analysis using PCA

The distance tree depicts the hierarchical clustering of all forty-six samples. Branch lengths represent the degree of similarity between dogs. The x-axis shows genetic distance, and branch values denote a confidence of 100. It highlights breed similarity by state where Punjab closely resembles Haryana and less with Karnataka (Figureure 1). In the Minimum Spanning Network (MSN) produced from reference-based SNP data, breed distinctions across states are visible through colours. The Punjab sample (Node Id 76 = Labrador retriever) at the centre likely represents ancestral forms and common SNPs. The branches radiate from the centre based on possible distances. Four elongated branches (Lab 21, Pug 44, Belgian Malinois 90, Lab 79) emerge from the same centre, indicating similarity within Punjab, Haryana, and Karnataka (Figureure 2).

**Figure 1:**
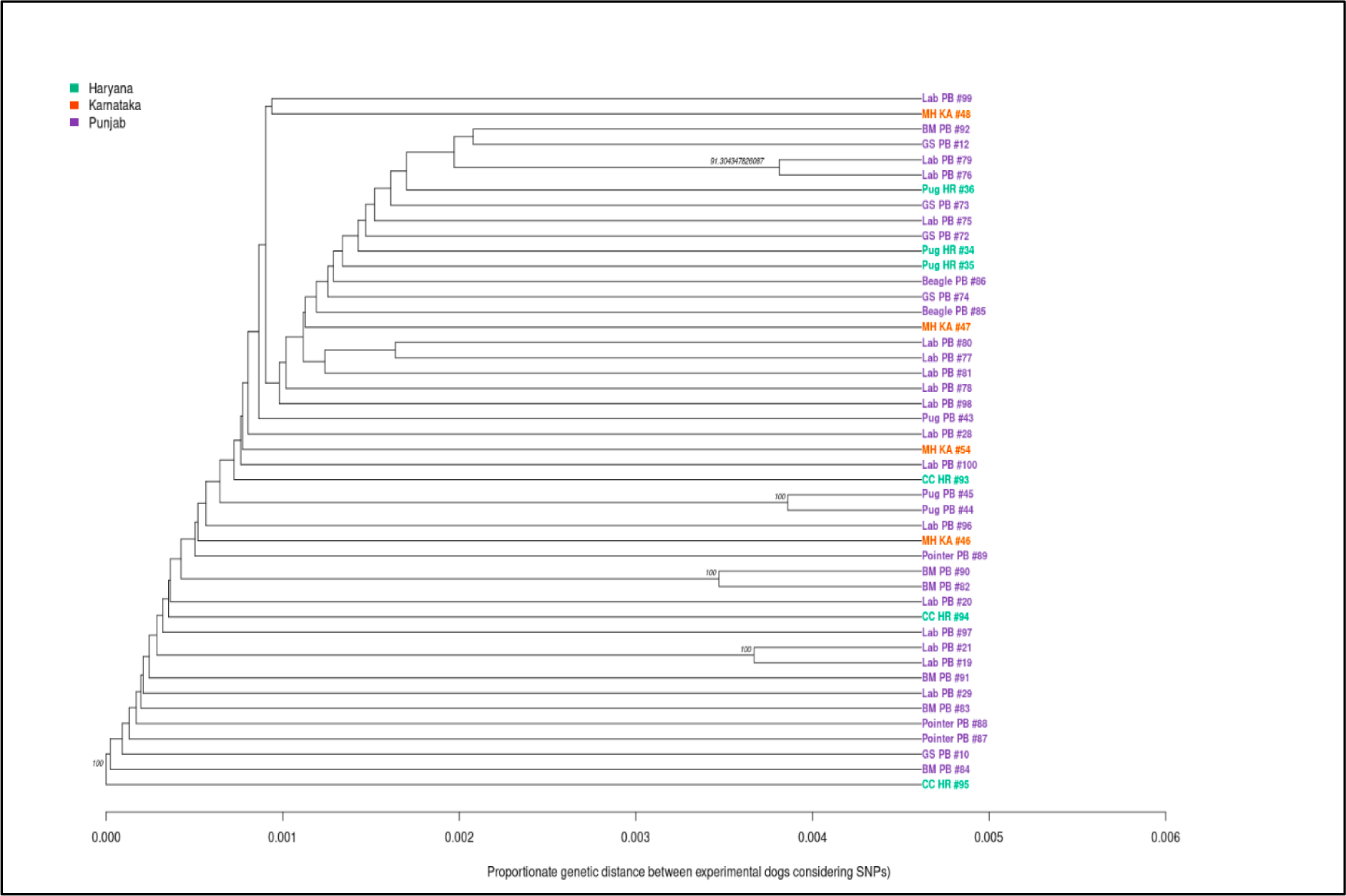
Distance tree indicating the genetic distance between the experimental groups with respect to location of sample collection.

**Figure 2:**
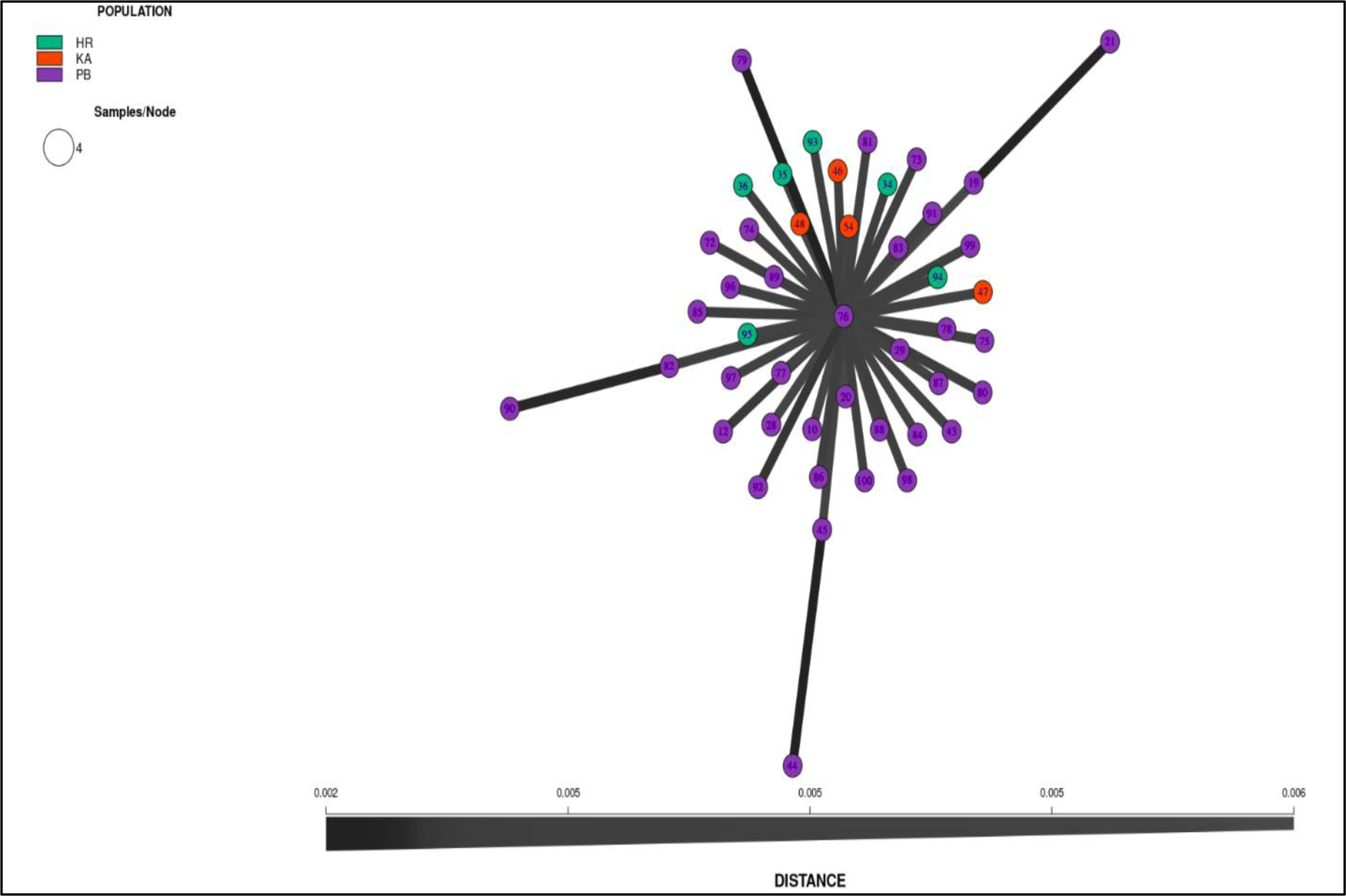
Minimum spanning network (MSN) between the experimental groups (eight divergent dog breeds)

PCA analysis revealed that PC2 distinguish Punjab (PB) and Karnataka (KA) samples (upper) from a cluster mainly comprising Punjab (PB) and Haryana (HR) samples (lower). Karnataka (KA) samples form a tightly clustered orange ellipse (Figureure 3). This pattern aligns with previous findings. Discriminant Analysis of Principal Components (DAPC) was used to delve into population assignments. DAPC situates breeds (dots) and states (three colours and ellipses) on a plane (Figureure 4). Analysing data using discriminant analysis (DA) aims to condense genetic differentiation. DAPC recognizes breed contributions to state clusters (Punjab, Haryana, Karnataka), aiding in identifying regions causing genetic divergence. PCA and DAPC outcomes are strikingly similar.

**Figure 3:**
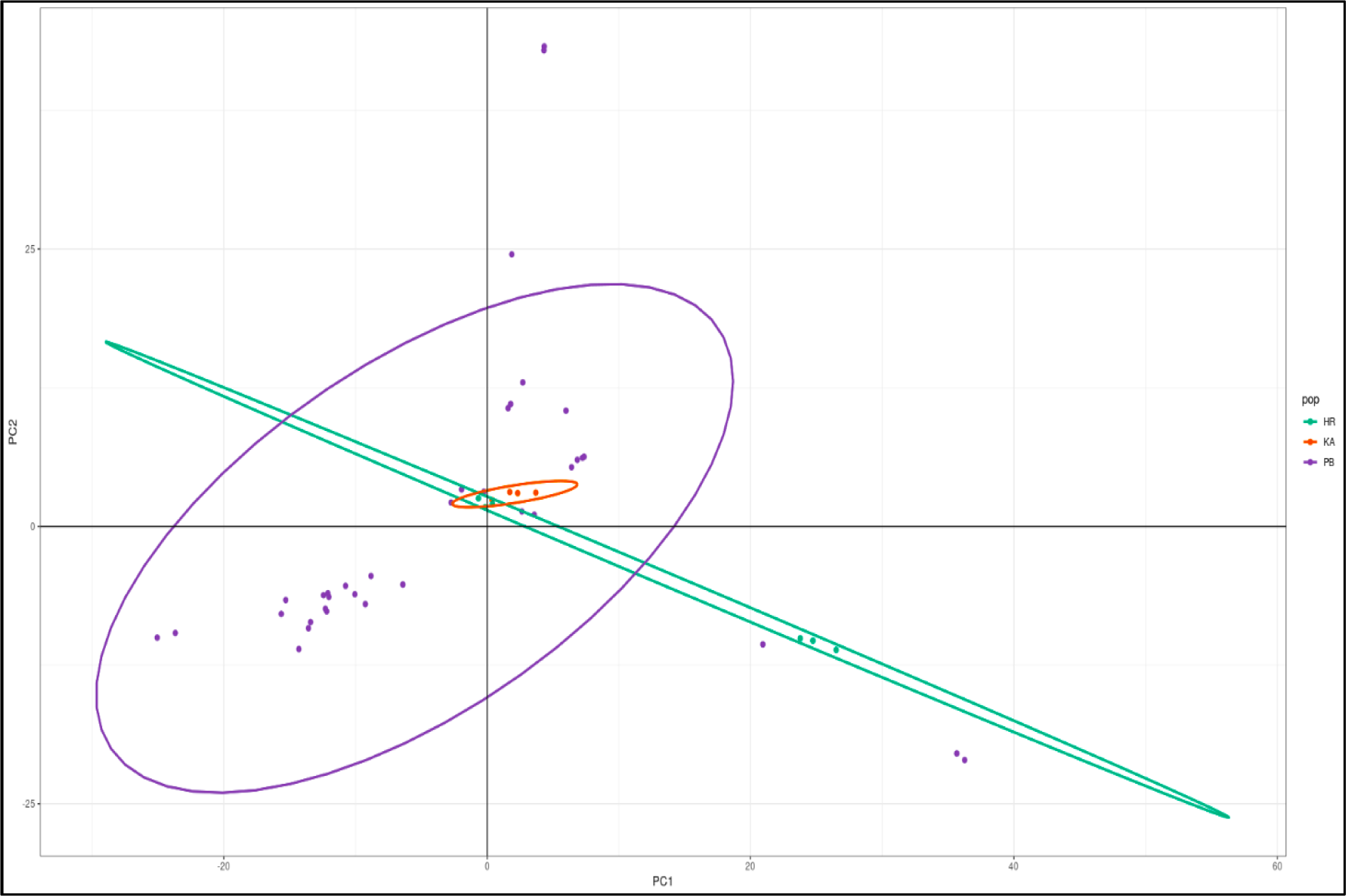
Principle components analysis (PCA) of divergent dog breeds.

**Figure 4:**
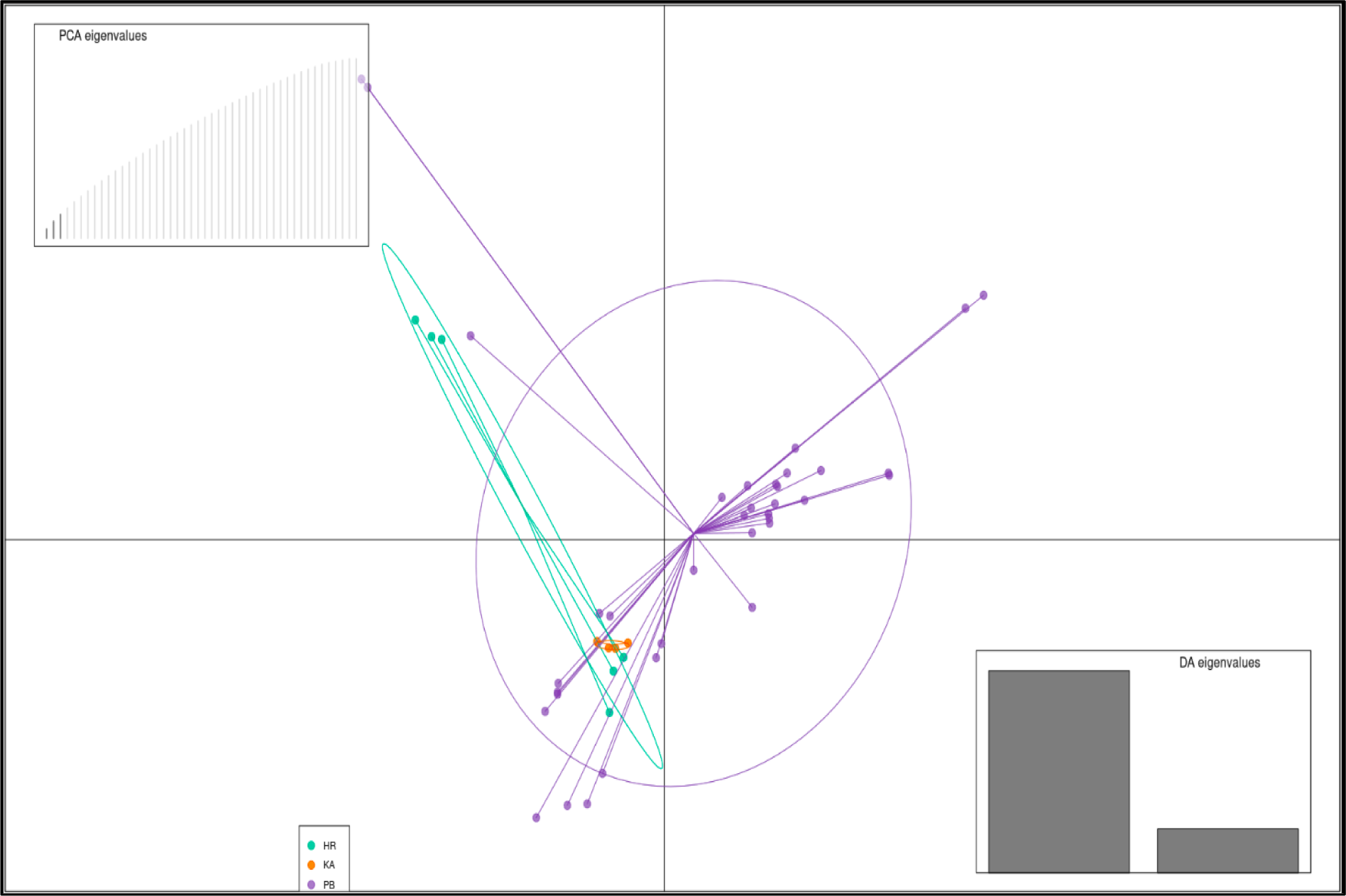
Discriminant analysis of principle components (DAPC) of divergent samples.

Membership probability determines the relatedness of each sample across geographical regions. It indicates the likelihood of a sample belonging to each state, visualized with coloured vertical lines (Figureure 5). Posterior probability refines this by updating estimates with new data. Distinct breeds sharing the same colour in the analysis indicate that they have a shared state of origin. This suggests that these breeds are genetically related and likely share common ancestry. Posterior membership probability categorizes samples by state and can be used to divide the sample into different states or regions. Each vertical line in the analysis represents an individual dog and they are separated based on their breed origin (Figureure 6). Plots reveal genetic diversity within populations. Breeds from Punjab resemble Haryana and Karnataka most, indicating close ancestral relationships and shared SNP genotypes. This suggests common ancestry between these breeds.

**Figure 5:**
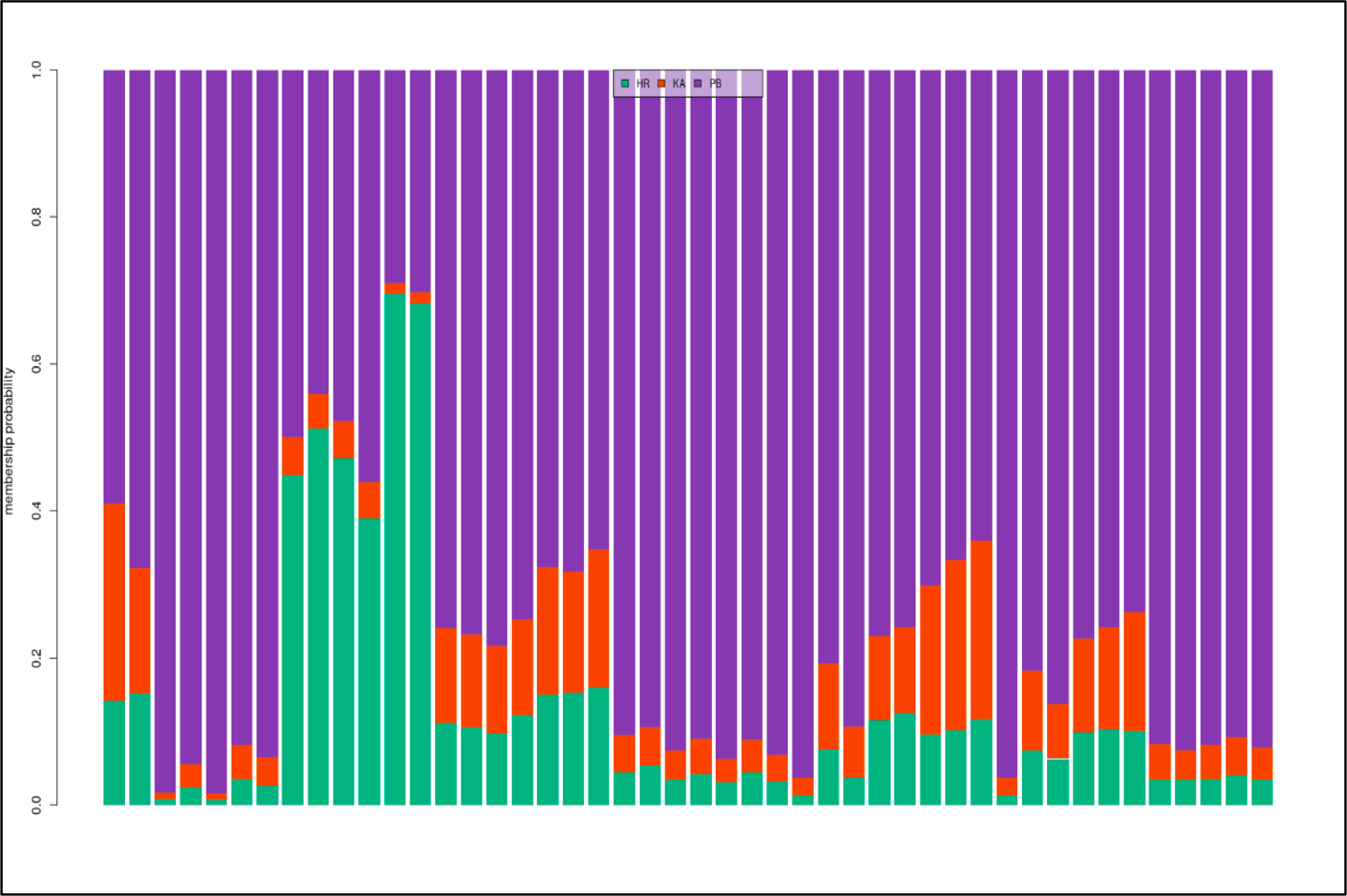
Membership probability of divergent samples.

**Figure 6:**
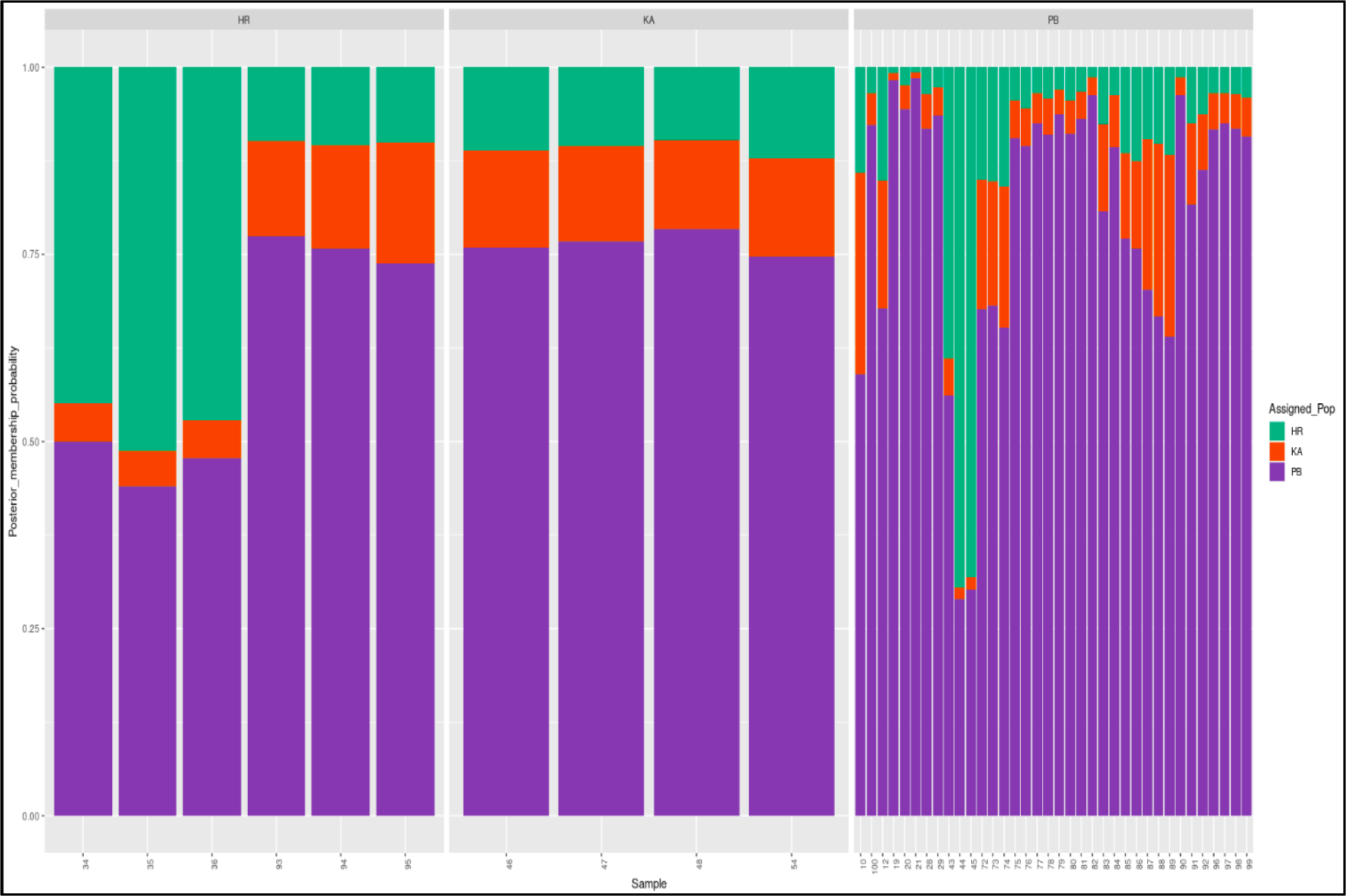
Posterior membership probability of divergent samples.

### Haplotype identification

Haplotype identification involves the identification of genetic variants that are located together on a chromosome, which was initially analyzed using R programming. The criteria used to determine haplotypes include, checking if the difference between SNP positions exceeded 80,000 (80kb) or if the cumulative sum of differences from the start of a haplotype surpassed 1,50,000 (150kb). When these conditions were met, and the number of SNPs within a haplotype was at least 4, the positions of the SNPs were stored in the ’haplotype_snps’ variable, and the haplotypes were added to the ‘haplotype’ list. Applying these criteria resulted in the identification of a total of 15,552 haplotypes out of the 2,18,429 SNP positions analyzed.

### Isolation and quality check of genomic DNA for haplotype validation

Blood samples from twenty-one different dogs were aseptically collected as mentioned earlier in Table 3. Genomic DNA was isolated using the standard Phenol: Chloroform: Isoamyl Alcohol (25:24:1) method (Green & Sambrook, 2018). DNA was then dissolved in TE buffer and stored at −20°C. DNA quality was confirmed by gel electrophoresis on a 0.8% Agarose gel (70V for 45-50 min.), with results observed on a UV Transilluminator. All samples displayed clear high molecular weight bands, deemed suitable for PCR amplification. DNA quantity was assessed by spectrophotometer, measuring optical density (OD) at 260/280 nm. Readings between 1.6 and 1.8 indicated good-quality DNA with no contamination.

### Identification of polymorphism

For the detection of SNPs, the reference sequence of respective chromosomes of *GBGT1*, *CELSR1* and *AFAP1* genes was aligned with the FASTA sequences received after sequencing, using Clustal Omega. In the present study, overall eleven SNPs were identified within each *GBGT1* (Table 4) and *CELSR1* (Table 5) genes, whereas no SNP was detected in the *AFAP1* gene. For SNP genotyping all chromatograms were screened manually for eleven detected SNP positions.

**Table 4.**
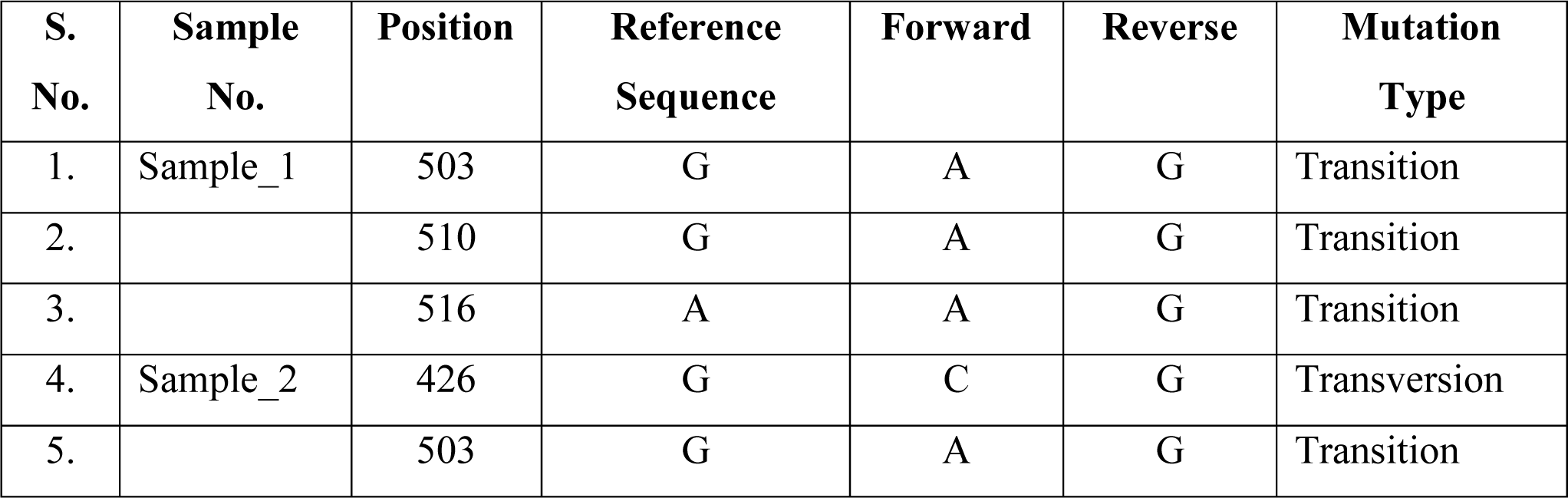
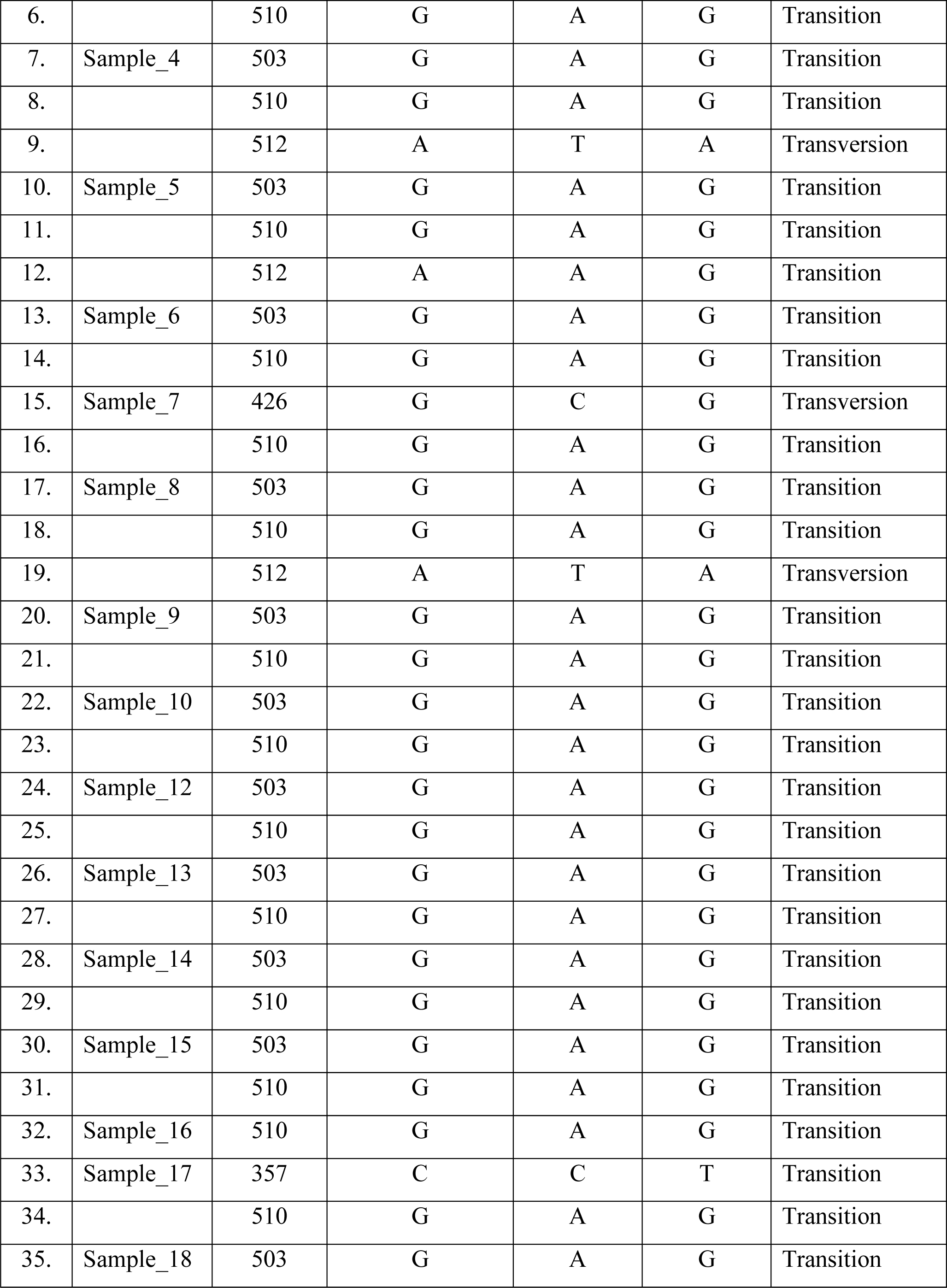
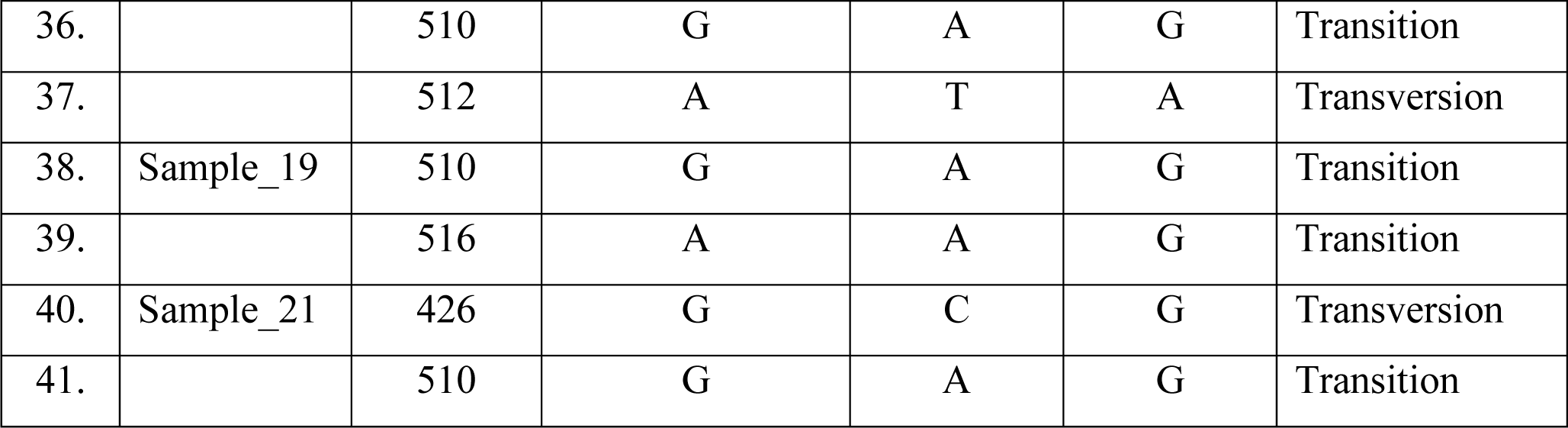
Summary of SNPs detected in the target genomic region of canine *GBGT1* gene.

The distribution of a total of five SNPs (426g>C, 503g>A, 510g>A, 512a>T and 516a>G) within the targeted region of canine *GBGT1* gene in different dog breeds. Among the observed SNPs, 503g>A was detected in samples 1, 2, 4, 5, 6, 8, 9, 10, 12, 13, 14, 15 and 18. At position 510, the SNP 510g>A was found in all the samples except in 3, 11 and 20. Only two samples, specifically 1 and 19 displayed the SNP 516a>G, which was absent in the remaining samples. The SNP 426g>C was exclusive to three samples viz. 2, 7 and 21. Sample numbers 4, 5, 8, and 18 exhibited the SNP 512a>T.

By employing the *CELSR1*-specific primers, a total of six SNPs (414g<A, 439g>A, 445 g>A, 469 g>A, 488 g>A and 667g<A) were identified in different dog breeds under study. The SNP 469 g>A was detected in samples 4, 7, 8, and 13. Also, the SNP 488g<A was found in 4, 7, and 8. Additionally, the SNPs 414g<A, 439g>A, 445 g>A and 469 g>A were detected in samples 4 and 7, whereas only SNP 667g<A was exclusively present in sample 7.

**Table 5.** Summary of SNPs detected in the target genomic region of canine *CELSR1* gene.

| S. No. | Sample No. | Position | Reference Sequence | Forward | Reverse | Mutation Type |
| --- | --- | --- | --- | --- | --- | --- |
| 1. | Sample_4 | 414 | G | A | G | Transition |
| 2. |  | 439 | G | A | G | Transition |
| 3. |  | 445 | G | A | G | Transition |
| 5. |  | 469 | G | A | G | Transition |
| 6. |  | 488 | G | A | G | Transition |
| 7. | Sample_7 | 414 | G | A | G | Transition |
| 8. |  | 439 | G | A | G | Transition |
| 9. |  | 445 | G | A | G | Transition |
| 10. |  | 469 | G | A | G | Transition |
| 11. |  | 488 | G | A | G | Transition |
| 12. |  | 667 | G | A | G | Transition |
| 13. | Sample_8 | 469 | G | A | G | Transition |
| 14. |  | 488 | G | A | G | Transition |
| 15. | Sample_13 | 469 | G | A | G | Transition |

### Pathway and enrichment analysis

The bioinformatics analysis of *AFAP1*, *CELSR1* and *GBGT1* genes carried out using the online David bioinformatics tool revealed the Glycosphingolipid biosynthesis – Globo and Isogloboseries pathway related to *GBGT1* gene in canines (Figureure 7) along with other genes (Table 6) involved. The *GBGT1* gene catalyzes the formation of Forssman glycolipid via the addition of GalNAc in alpha-1,3-linkage to Gb4Cer. Forssman glycolipid (also called Forssman antigen; FG) serves for adherence of some pathogens such as *E.coli* uropathogenic strains. These uropathogenic strains of *E.coli* are the most common cause of urinary tract infections in dogs (Yousefi & Torkan, 2017).

**Figure 7:**
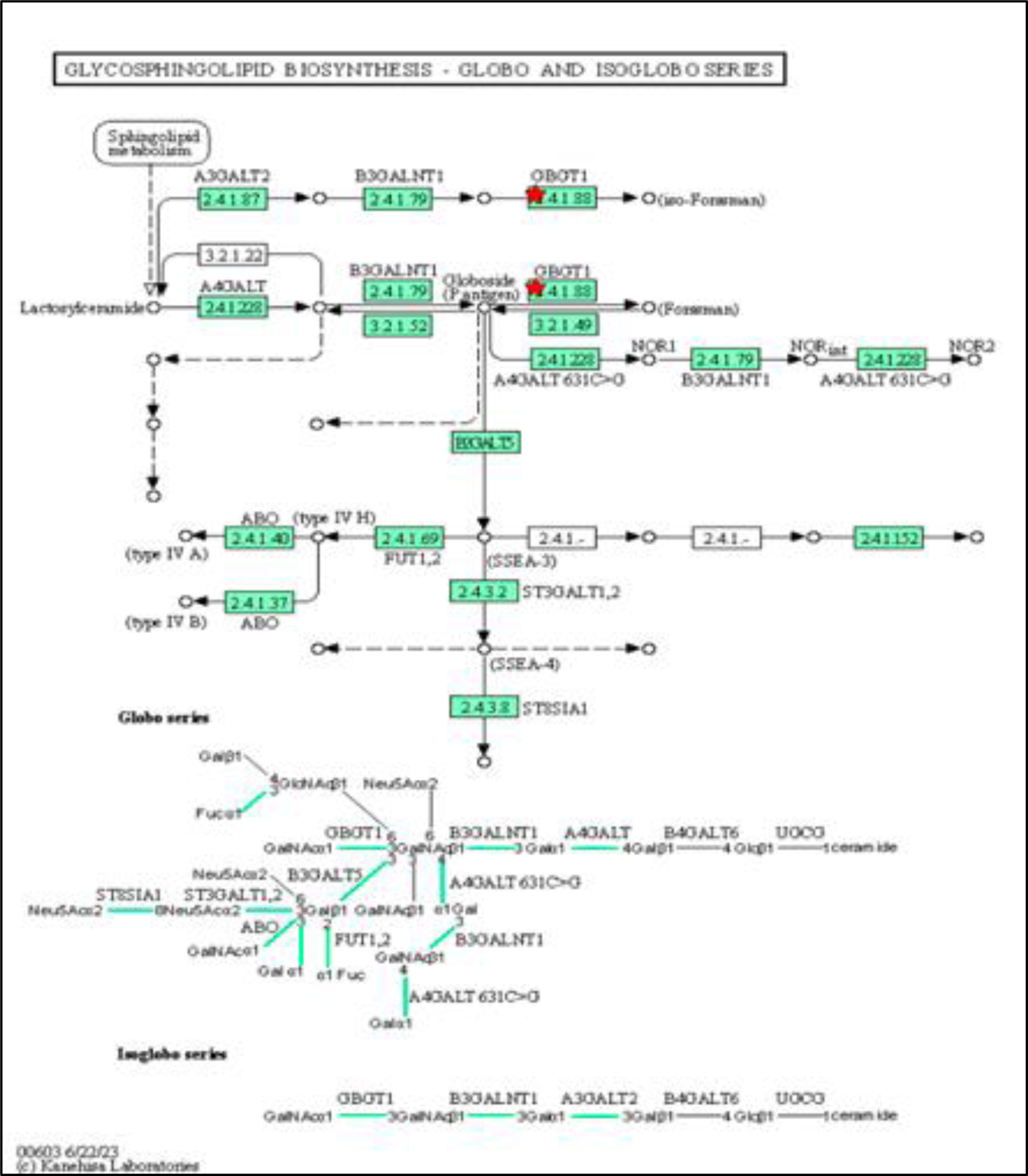
KEGG_PATHWAY result for *GBGT1* gene showed Glycosphingolipid biosynthesis – Globo and Isogloboseries.

**Table 6.**
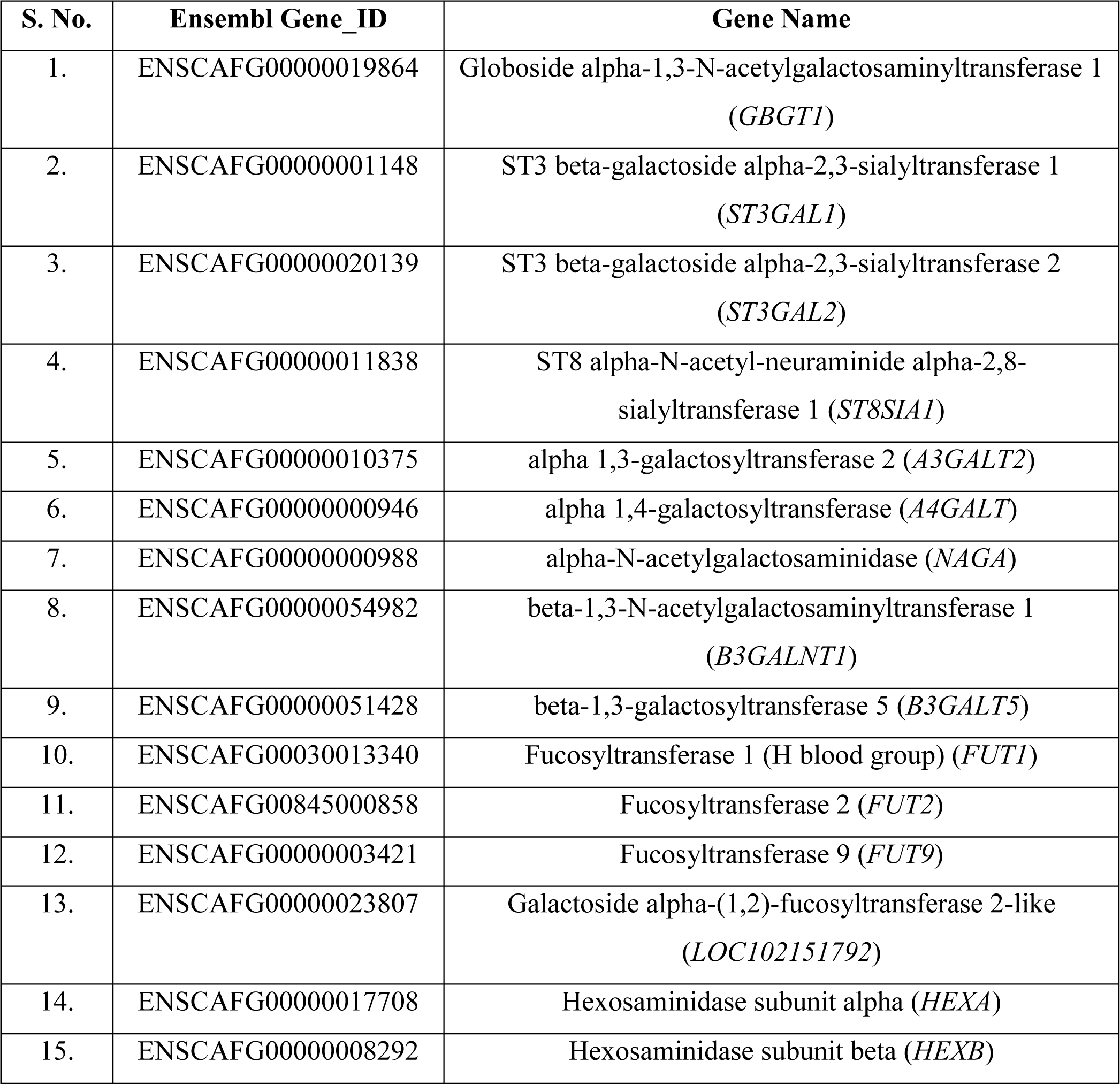
Details of the genes involved in Glycosphingolipid biosynthesis – Globo and Isogloboseries pathway related to *GBGT1* gene with corresponding Ensembl gene names.

The three genes were subjected to the PANTHER 17.0 tool for analyzing the biological process, molecular function and pathway analysis. The results show the list of three genes with their specific gene ID, gene family, symbols, and their mapped ID. The functional classification of all the genes with the gene’s name, genes IDs, Ensembl IDs, and different classes of proteins showed in which they belong. The pie chart (Figureure 8) shows the functional classification of the genes. The results of PANTHER classification by molecular function analysis of three genes with associated gene ontology (GO) terms. Binding activities GO:0005488 (red), Catalytic activities GO:0003824 (blue) and No PANTHER category is assigned (UNCLASSIFIED) (violet). Under the biological function (Figureure 9) a total of four hits were generated, out of which for No PANTHER category is assigned (UNCLASSIFIED) (violet), Biological adhesion GO:0022610 (red), Cellular process GO:0009987 (blue) and Metabolic process GO:0008152 (green). A total of three genes were subjected to pathway analysis, only two pathways i.e., Cadherin signaling pathway (P00012) and Wnt signaling pathway (P00057) for the *CELSR1* gene were found (Figureure 10).

**Figure 8:**
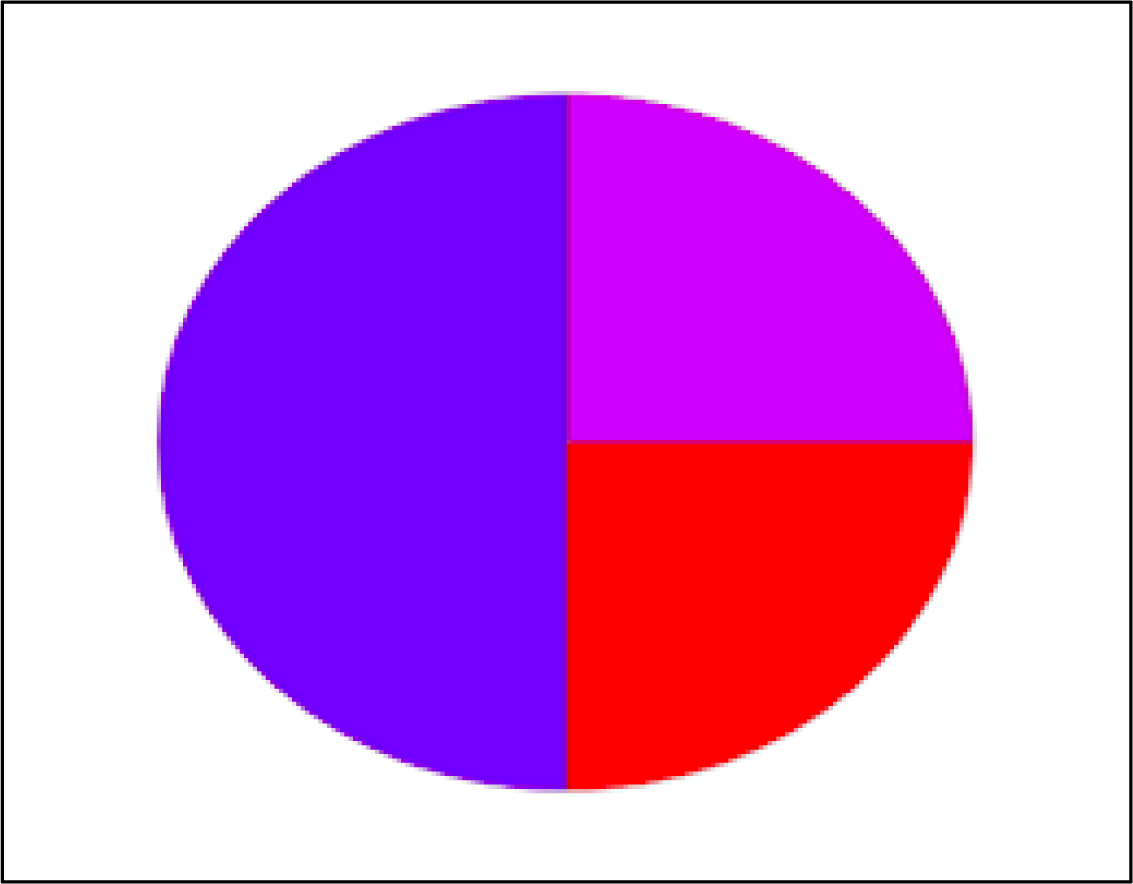
Molecular functions of *AFAP1*, *CELSR1* and *GBGT1* genes.

**Figure 9:**
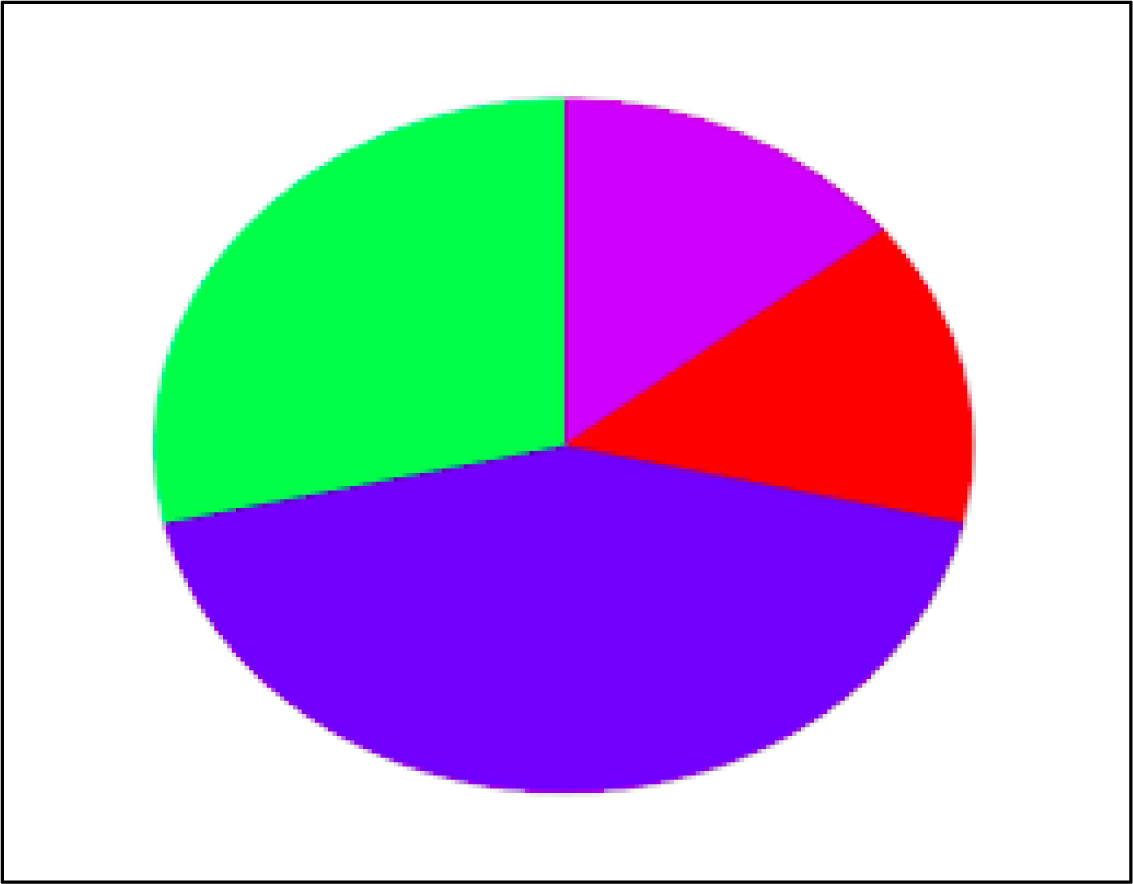
Biological process of *AFAP1*, *CELSR1* and *GBGT1* genes.

**Figure 10:**
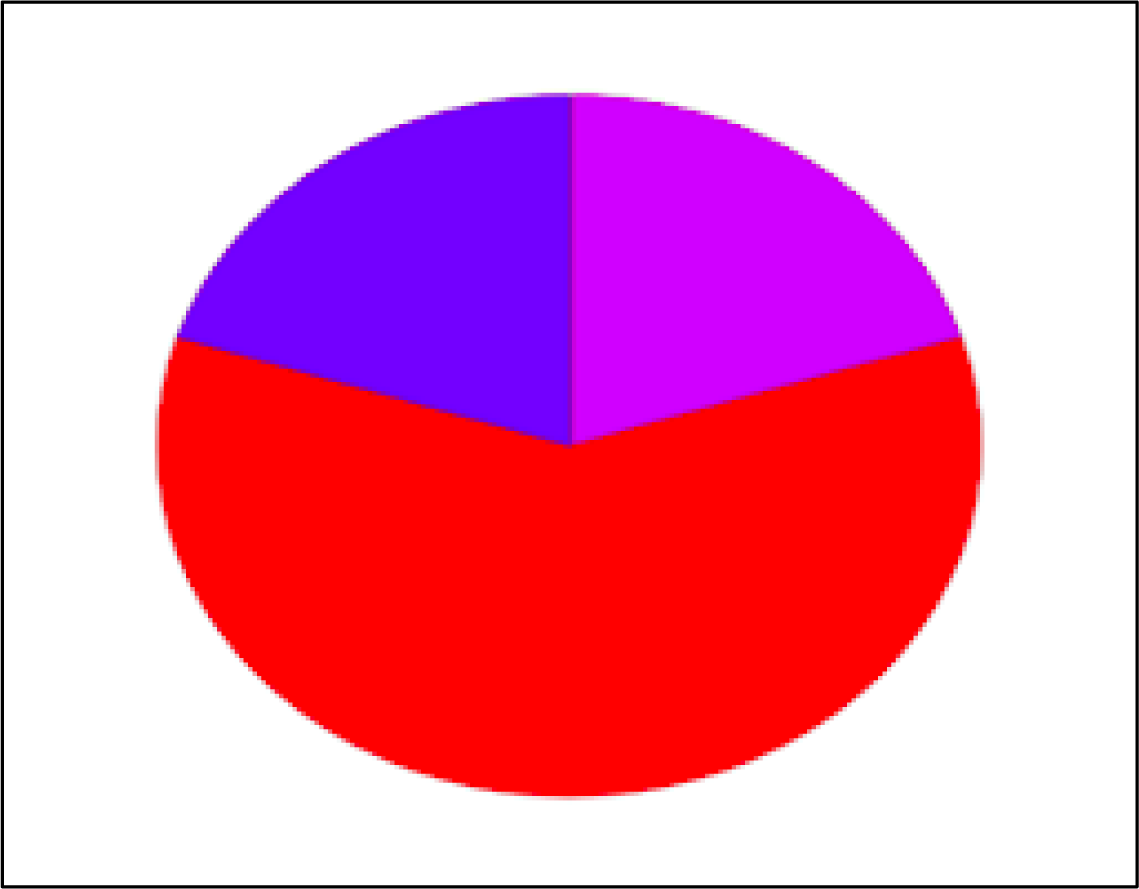
Pathway analysis of *AFAP1*, *CELSR1* and *GBGT1* genes.

GeneCards version 5.12 is significantly used for understanding complex systems biology. It offers a platform for the association of gene expression, GO ID, function and pathways. All the genes were studied for their function in humans and were extracted in terms of Gene Ontology (GO) ID and the pathways associated with each gene. The details of each gene are shown below in Table 7.

**Table 7.** Pathway analysis of genes using GeneCards version 5.12.

| S. No. | Gene Name | Pathways | Functions | GO ID | GO terms |
| --- | --- | --- | --- | --- | --- |
| 1. | Actin Filament Associated Protein 1 (*AFAP1*) | Immune response_IFN gamma signaling pathway | May Modulate changes in actin filament integrity and induce lamellipodia formation | GO:0003779 3 5 | actin binding |
|  |  |  |  | GO:0017124 3 5 | SH3 domain binding |
|  |  |  |  | GO:0042169 3 5 | SH2 domain binding |
| 2. | Cadherin EGF LAG Seven-pass G-type Receptor 1 (*CELSR1*) | G Protein Coupled Receptors (GPCRs) | Receptor molecule that may have an important role in cell/cell signalling during nervous system formation | GO:0004888 5 | transmembrane signaling receptor activity |
|  |  |  |  | GO:0004930 3 5 | G protein-coupled receptor activity |
|  |  |  |  | GO:0005509 3 5 | calcium ion binding |
| 3. | Globoside alpha-1,3-N-acetylgalactosaminyltransferase 1 (*GBGT1*) | Superpathway of glycosphingolipids biosynthesis | Has lost the ability to synthesize Forssman glycolipid antigen and might have acquired an alternative function in glycosphingolipid metabolism | GO:0016740 5 | transferase activity |
|  |  |  |  | GO:0016757 5 | glycosyltransferase activity |
|  |  |  |  | GO:0016758 5 | hexosyltransferase activity |
|  |  |  |  | GO:0046872 3 5 | metal ion binding |
|  |  |  |  | GO:0047277 3 5 | globoside alpha-N-acetylgalactosaminyltransferase activity |

### Evolutionary relationships between dogs under study: Phylogenetic tree construction and pairwise distance matrix

Phylogenetic analysis of the twelve obtained sequences of *AFAP1* (AF3A & AF3B fragments), *CELSR1* and *GBGT1* for Labrador Retriever and German Shepherd dogs were constructed individually. It revealed different clusters for each phylogenetic tree (Figureures 11, 12, 13 & 14) which depicts that there exists genetic differentiation between the samples. Each cluster contains a group of samples with shared SNPs or genetic history. The formation of clusters signifies that there exists a population structure among the samples used for validation studies. Furthermore, pairwise distance matrix validates the cluster formation by computing exact genetic distances between all pairs of samples for *AFAP1* AF3A fragment (Figureure 15), *AFAP1* AF3B fragment (Figureure 16), *CELSR1* gene (Figureure 17) and *GBGT1* gene (Figureure 18).

**Figure 11:**
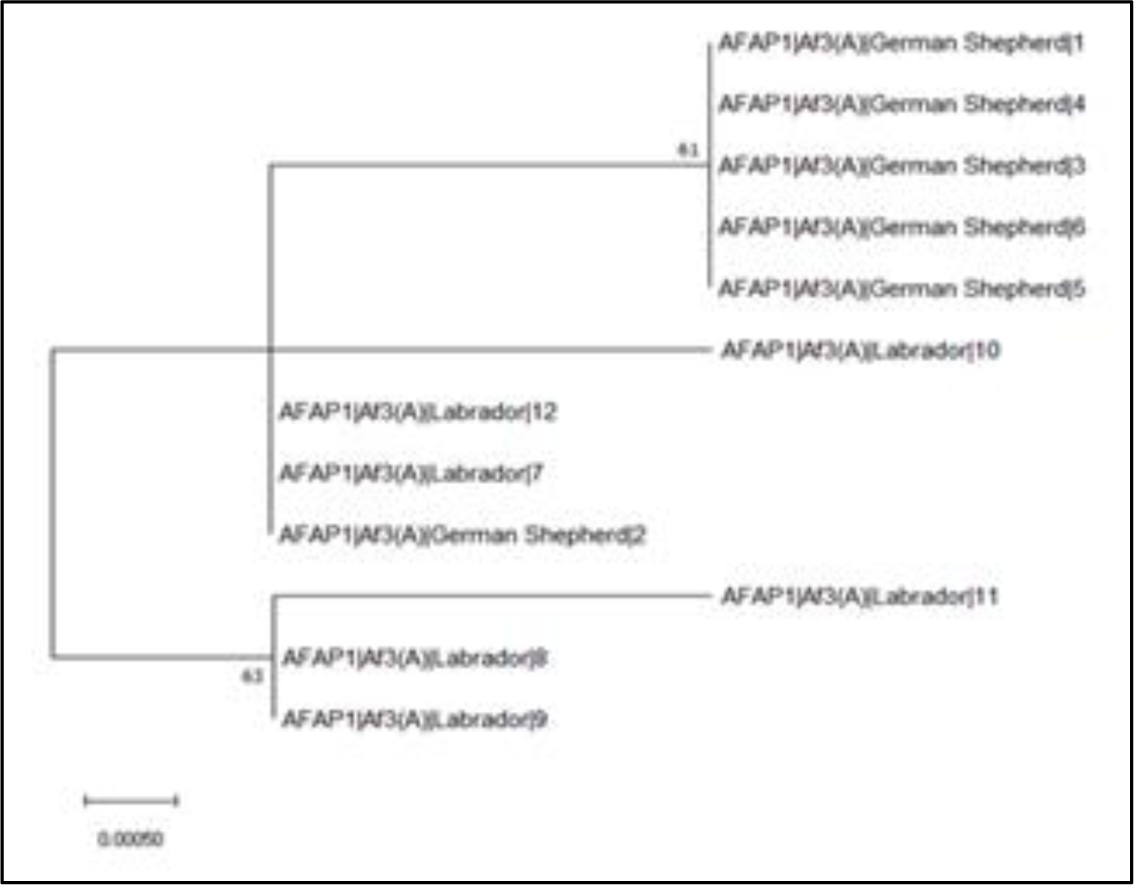
Phylogenetic tree constructed by utilizing twelve FASTA sequences for *AFAP1* gene AF3A fragment using maximum likelihood model with 1000 bootstraps resampling using MEGA 11 software.

**Figure 12:**
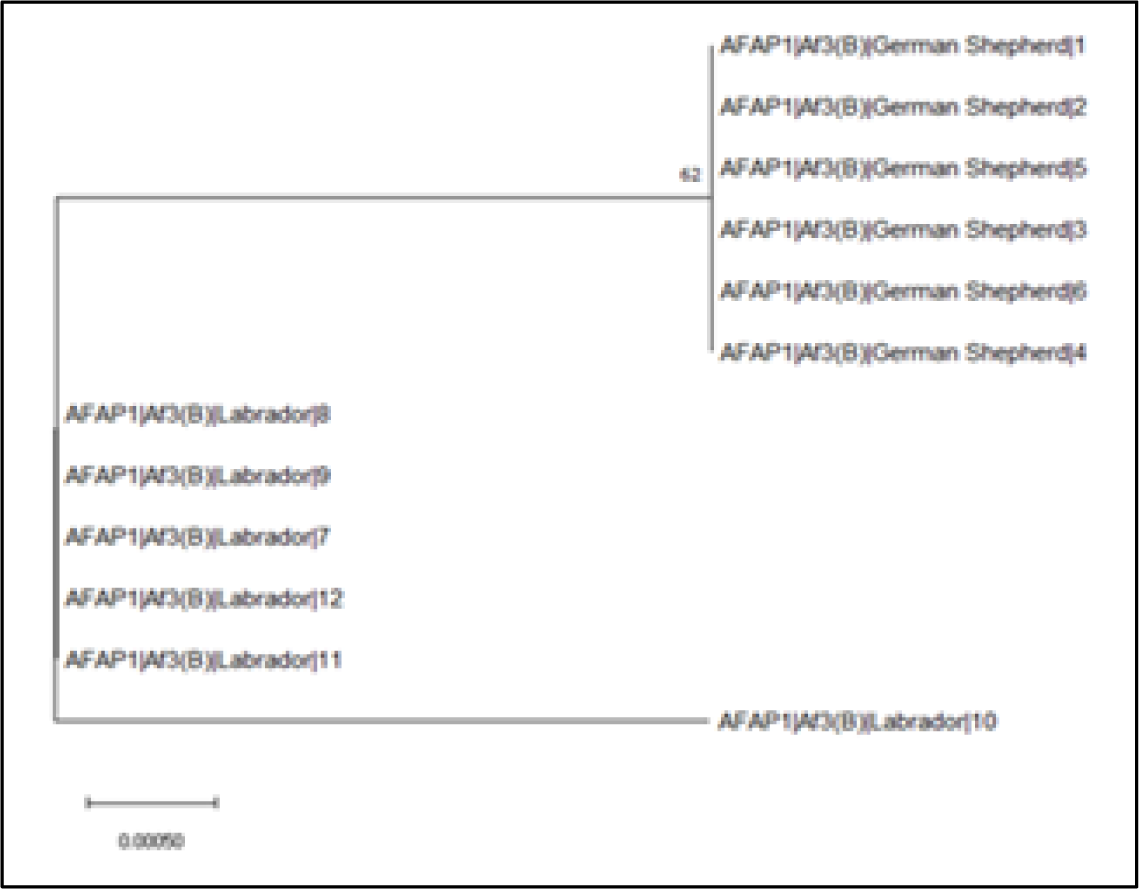
Phylogenetic tree constructed by utilizing twelve FASTA sequences for *AFAP1* gene AF3B fragment using maximum likelihood model with 1000 bootstraps resampling using MEGA 11 software.

**Figure 13:**
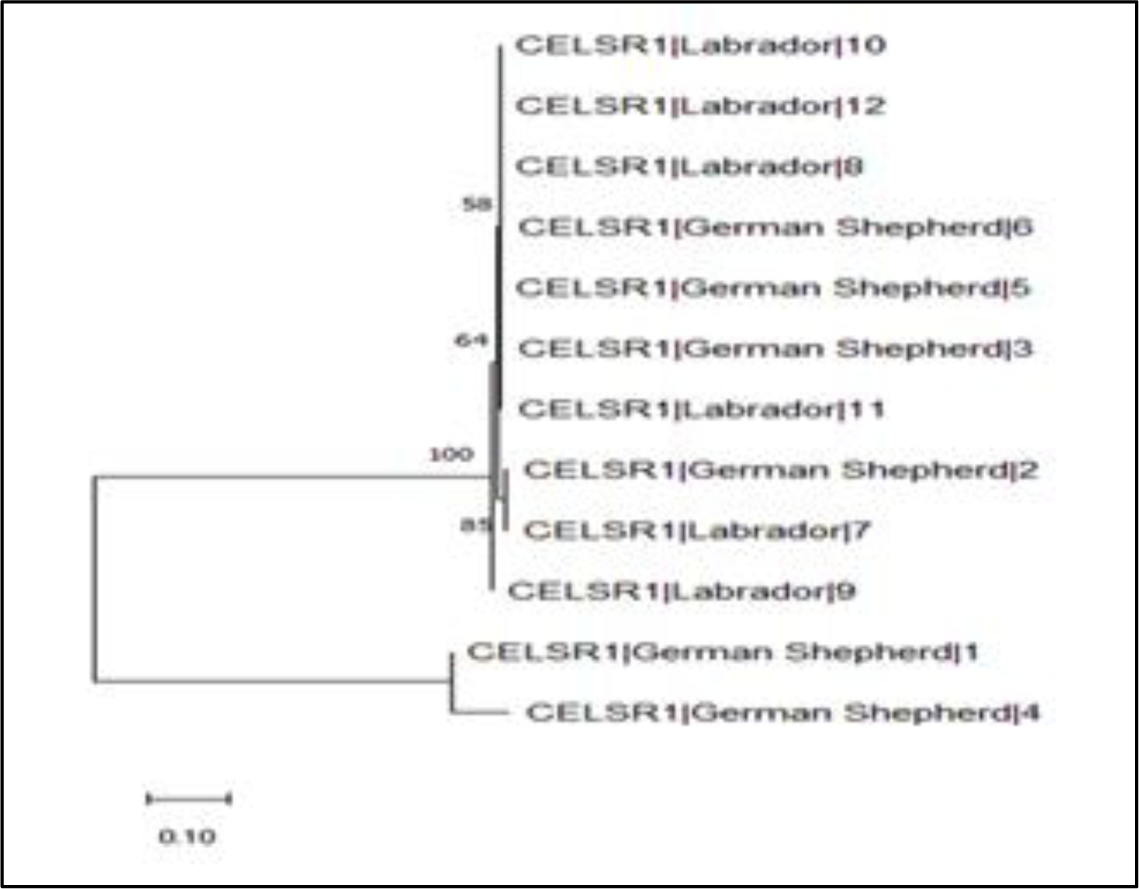
Phylogenetic tree constructed by utilizing twelve FASTA sequences for *CELSR1* gene using maximum likelihood model with 1000 bootstraps resampling using MEGA 11 software.

**Figure 14:**
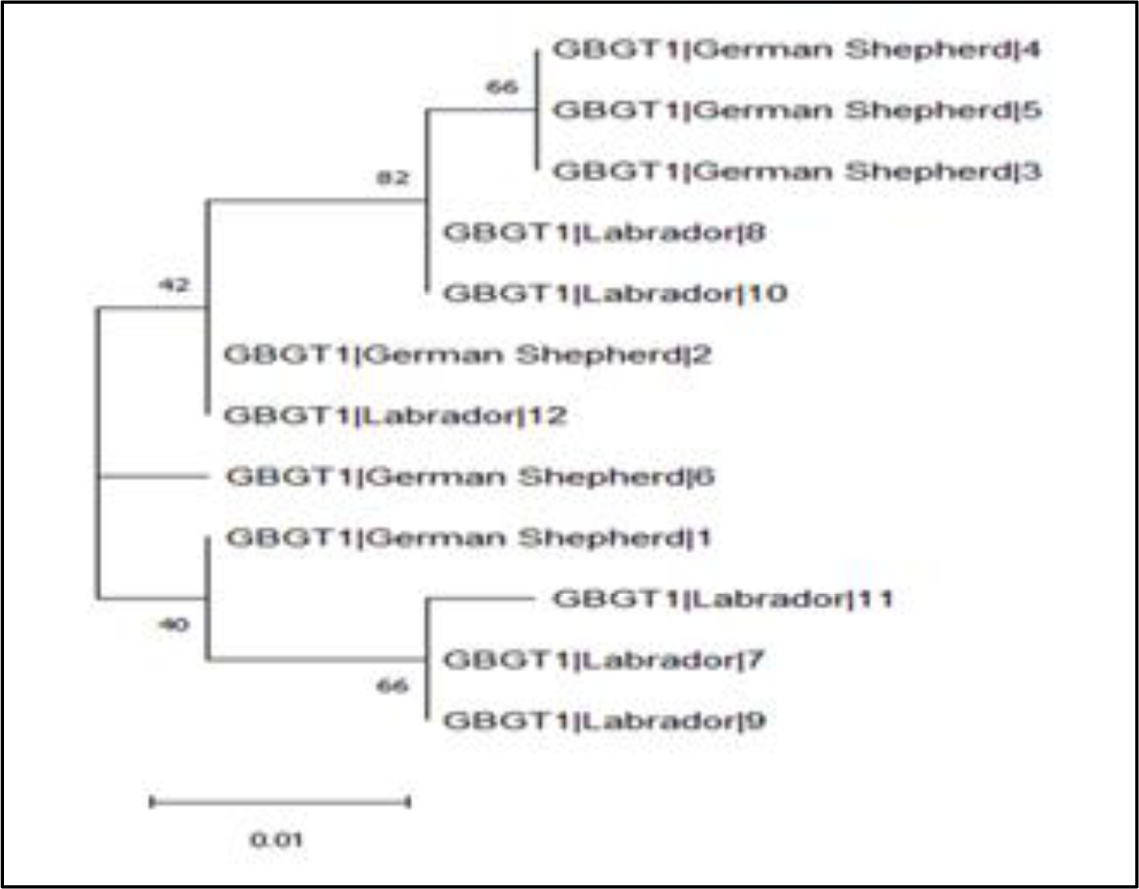
Phylogenetic tree constructed by utilizing twelve FASTA sequences for *GBGT1* gene using maximum likelihood model with 1000 bootstraps resampling using MEGA 11 software.

**Figure 15:**
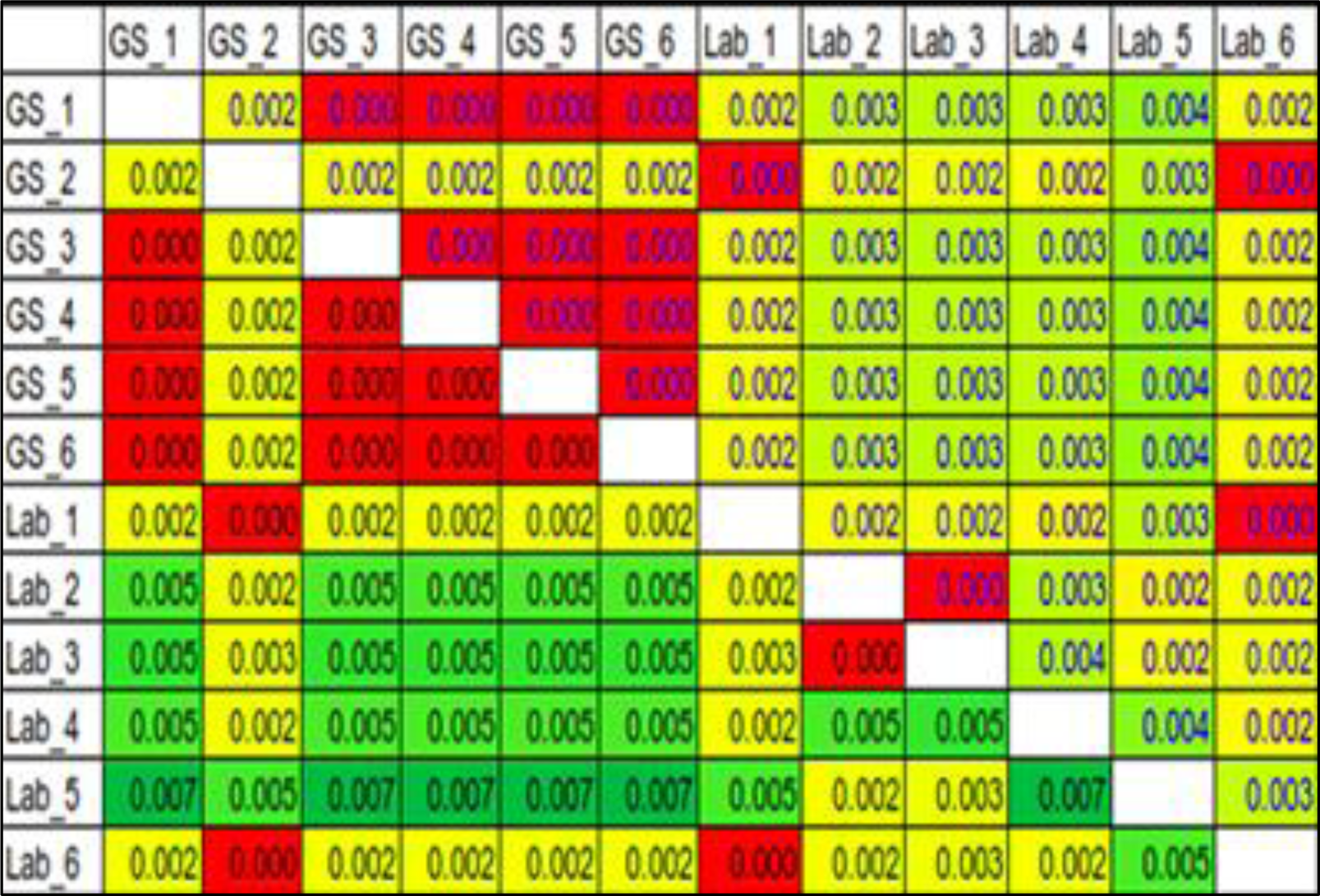
Pairwise distance matrix calculated by utilizing twelve FASTA sequences for *AFAP1* gene AF3A fragment using MEGA 11 software.

**Figure 16:**
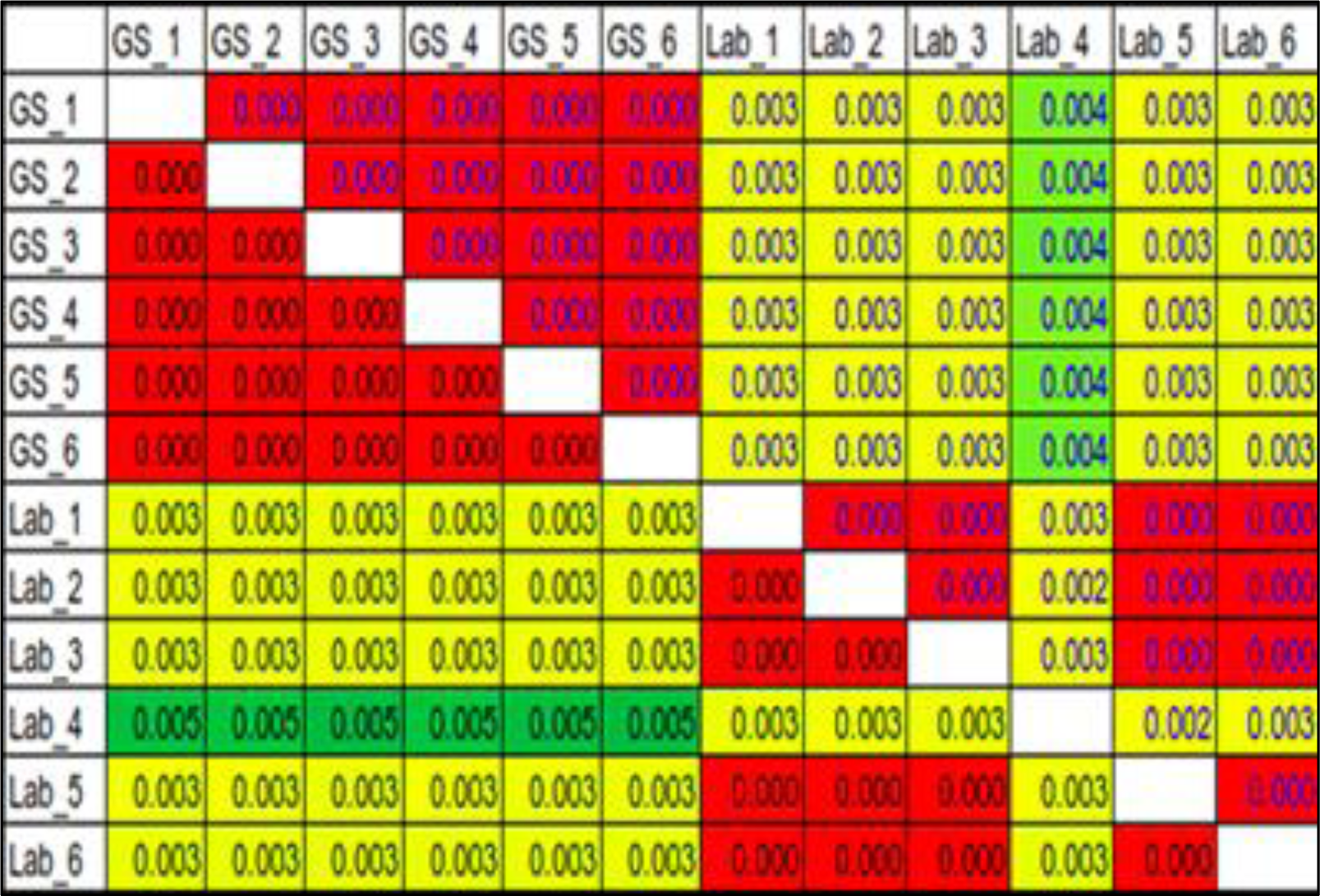
Pairwise distance matrix calculated by utilizing twelve FASTA sequences for *AFAP1* gene AF3B fragment using MEGA 11 software.

**Figure 17:**
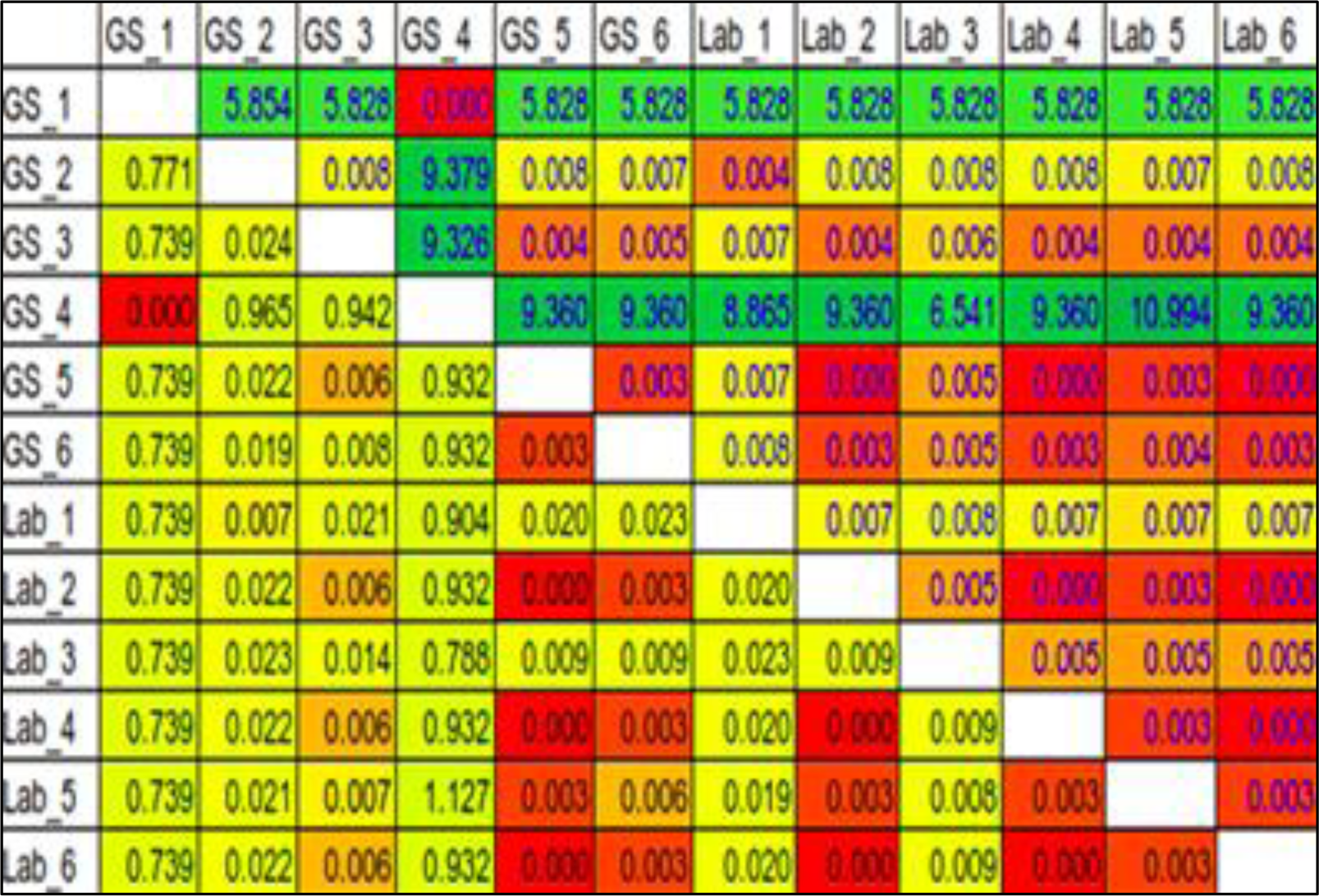
Pairwise distance matrix calculated by utilizing twelve FASTA sequences for *CELSR1* gene using MEGA 11 software.

**Figure 18:**
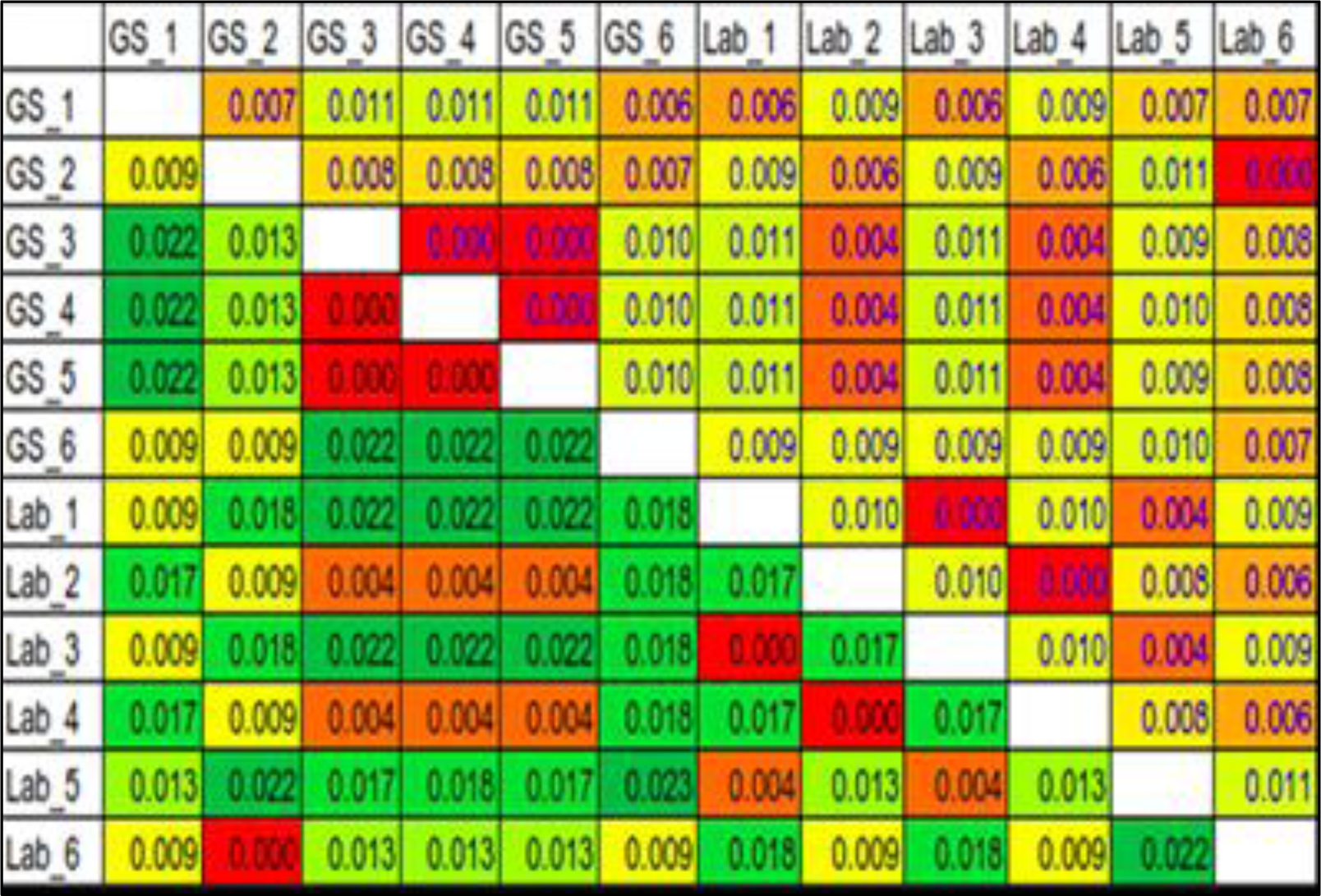
Pairwise distance matrix calculated by utilizing twelve FASTA sequences for *GBGT1* gene using MEGA 11 software.

## Discussion

Dog (*Canis lupus familiaris)* is a domesticated animal that descended from wolves. It is commonly referred to as the domestic dog and is believed to have originated from wolves that existed during the Pleistocene era. Dogs were the first species to be domesticated by hunter-gatherers more than 15,000 years ago, before the development of agriculture.

Despite the significant role that dogs play in our history, surprisingly little attention has been given to understanding the genetic makeup of dog breeds found in India. The specific variations (SNPs) in canine DNA unique to India remain largely unexplored. However, by investigating the population structure and identifying distinct haplotypes, these existing constraints can be overcome and new possibilities can be unlocked. Therefore, the research endeavours to unravel the genetic structure of prominent dog breeds found in both North and South India. This can be achieved by meticulously analysing a vast array of genome-wide single-nucleotide polymorphism (SNP) data generated through ddRAD technology using the Illumina 150 bp paired-end sequencing method.

Genotyping-by-sequencing (GBS) is one such approach to obtain high-frequency SNPs. The strategy has been used for population genetic studies, and association mapping, and has proven to be a powerful tool to dissect multiple genes/QTLs (Reddy *et al*. 2020). We obtained 8,13,580 unfiltered SNPs of which 2,18,433 high-quality SNPs were used for studying population structure in divergent dog breeds. Kaur *et al*. (2023) conducted a study to determine the genome-wide distributed SNPs and haplotypes in indigenous Gaddi dogs and other popular dog breeds maintained in India. They screened 3,56,461 SNPs loci, out of which 75,811 SNPs were used.

Distance tree and minimum spanning network (MSN) produce similar outcomes. Also, the patterns produced by principle component analysis (PCA) and discriminant analysis of principle components (DAPC) are highly similar and depict how similar the breeds are based on their geographical locations. All the samples collected from different states of India (Punjab, Haryana and Karnataka) suggest close ancestral relationships between breeds indicating that they share the common ancestral SNP genotypes. Furthermore, the presence of distinct clusters observed during phylogenetic tree construction using the maximum likelihood model, confirms the genetic differentiation between the samples used for validation studies. Each cluster demonstrated the group of samples with a shared genetic history and the formation of clusters shows that there exists a clear population structure among the samples. The pairwise distance matrix validates the cluster formation by computing exact genetic distances between all pairs of samples under study.

Depending on the species and population used, haplotype blocks made up of SNPs in significant LD can span up to a few Mb. The haplotypes defines the genetic variation across the genome. Conserved haplotypes can then be used for the identification and characterization of functionally important genomic regions during evolution and/or selection (Qian *et al*. 2014). In this study, we have identified 15,552 haplotypes using SNPs produced from ddRAD-GBS. From the total haplotypes identified, three haplotypes were selected based on the number of SNPs, the haplotypes having more SNPs have a greater chance of capturing the genetic variation, and the SNP with a minor allele frequency (MAF) greater than 0.05 or 5% were targeted.

Out of the three chosen haplotypes, haplotype-1 is located on chromosome 3, spanning a length of 187bp and containing four SNPs, and was found to be linked with the *AFAP1* gene. Regarding haplotypes 2 and 3, they have lengths of 111bp and 117bp, positioned on chromosomes 9 and 10. They encompass five and six SNPs each and are associated with the *GBGT1* and *CELSR1* genes respectively. The customized Sanger sequencing results revealed a total of nine SNPs, five present in the *GBGT1* gene and the remaining four in the *CELSR1* gene. However, no SNP was found in the *AFAP1* gene as there were no evident peaks present in the electrophoregrams. These three genes were further analyzed using DAVID, PANTHER and GeneCards database for pathway enrichment analysis, molecular function, cellular function, biological function and gene expression.

From the pathway analysis, the Glycosphingolipid biosynthesis – Globo and isogloboseries pathway linked with the *GBGT1* gene in dogs was found. The *GBGT1* gene in dogs catalyzes the formation of Forssman glycolipid via the addition of GalNAc in alpha-1,3-linkage to Gb4Cer (Haslam & Baenziger, 1996). Forssman glycolipid (also called Forssman antigen; FG) probably serves for adherence of some pathogens such as *E.coli* uropathogenic strains (Xu *et al*. 1999). These uropathogenic strains of *E.coli* are the most common cause of urinary tract infections in dogs (Yousefi & Torkan, 2017). Contradicting this, Jesus *et al*. (2018) proved that due to the presence of two missense mutations, *GBGT1* is a pseudogene in humans. Instead, humans develop the naturally occurring anti-Forssman antibody, which, when interacting with the Forssman-antigen, activates the complement system.

In gene ontology (GO) analysis, relevant pathways for *CELSR1* genes in dogs are the ‘Cadherin signaling pathway’ and ‘Wnt/PKC pathway’. The Cadherin signaling pathway involves calcium ion-dependent adhesion facilitated by the cadherin gene family, including E-cadherin, N-cadherin, and P-cadherin. The Cadherin-catenin complex is crucial, affecting processes like development, neurogenesis, adhesion, and inflammation. It is linked to diseases like cancer (Kourtidis *et al*. 2017).

The Wnt pathway starts with Wnt binding to Frizzled receptors on neighbouring cells, leading to dishevelled protein activation and an unknown G protein activation. This inhibits the Beta-Catenin destruction complex, increasing cytosolic Beta-Catenin. This excess beta-Catein moves to the nucleus and binds with transcriptional regulatory molecules like TCF/LEF proteins, inducing gene transcription. Activation of the G-protein pathway prompts Phospholipase C activation which catalyzes the conversion of PI(4,5)P2 to DAG and IP3 (Kadamur & Ross, 2013). *CELSR1* has neuro-protective effects in cerebral ischemia (which is the acute brain injury that results from the impaired blood flow to the brain), by reducing cell apoptosis in the cerebral cortex and promoting neurogenesis and angiogenesis. Therefore, *CELSR1* can be a potential target for the clinical treatment of cerebral ischemia (Wang *et al*. 2020).

The role of the *AFAP1* gene in dogs remains unclear, but human gene expression studies from GeneCards indicate its involvement in modulating actin filament integrity and inducing lamellipodia formation. *AFAP1* interacts directly with actin filaments via a carboxy terminus actin-binding domain, suggesting its role as an adapter protein connecting actin filaments to Src family members and signaling proteins (Qian *et al*. 2002). The capacity of *AFAP1* to modify actin filament integrity is demonstrated by conformational changes induced by phosphorylation or mutagenesis (Dorfleutner *et al*. 2008). This suggests the potential roles of *AFAP1*, including acting as an adaptor protein for signaling molecules, forming larger signaling complexes, activating Src family kinases due to conformational changes, and directly impacting actin filament organization as a cross-linking protein.

## Conclusions

Our study focused on analyzing the genetic makeup of popular canine breeds from both north and south India. This analysis provided new insights into the genetic composition, specific genetic variations (SNPs), and unique DNA sequences (haplotypes) of distinct Indian dog breeds. By studying the population structure, we revealed how different breeds are genetically connected based on their geographical locations. Through genotyping-by-sequencing (GBS) of the dogs under study, we identified a substantial number of 2,18,433 SNPs and 15,552 haplotypes. These haplotypes could potentially serve as markers for studying diseases and other traits. The diverse SNP variations we found also shed light on the genetic diversity of dog populations in India, which has valuable implications for dog breeders. Additionally, we conducted pathway enrichment analysis on the genes associated with the haplotypes, confirming their molecular and biological functions. Overall, our research contributes to the molecular characterization of indigenous dog breeds and offers potential insights for future studies and breed management.

## Funding

The authors thankfully acknowledge the funding provided by the Department of Biotechnology, Government of India, through the collaborative research project “Parentage Determination and Cytogenetic Profiling in Dogs (DBT-19I)”.

## Author contributions

DS: material collection, wet lab analysis for validation study, data analysis and manuscript writing; SKM: provided with the blood samples for validation studies; NK: manuscript editing; CSM: designing, supervising the work and manuscript editing. All authors read and approved the manuscript. All authors contributed to manuscript revision, read, and approved the submitted version.

## Acknowledgements

The authors are grateful to the University for providing research facilities and conducive environment for conducting the study. Sincere thanks to the sequencing agency (NovoGene, Mumbai) for providing the preliminary data analysis.

